# HCN1 channels mediate mu opioid receptor long-term depression at insular cortex inputs to the dorsal striatum

**DOI:** 10.1101/2021.08.31.458358

**Authors:** Braulio Munoz, Brandon M. Fritz, Fuqin Yin, Brady K. Atwood

**Author notes:** **Author Contact information Corresponding author**: Brady K. Atwood, PhD., Indiana University School of Medicine, Department of Pharmacology & Toxicology, 320 W 15th St., NB 400C, Indianapolis, IN, USA 46202.

## Abstract

Mu opioid receptors (MORs) are expressed in the dorsal striatum, a brain region that mediates goal-directed (via the dorsomedial striatum), and habitual (via the dorsolateral striatum, DLS) behaviors. Our previous work indicates that glutamate transmission is depressed when MORs are activated in the dorsal striatum, inducing MOR-mediated long-term synaptic depression (MOR-LTD) or short-term depression (MOR-STD), depending on the input. In the DLS, MOR-LTD is produced by MORs on anterior insular cortex (AIC) inputs and MOR-STD occurs at thalamic inputs, suggesting input-specific MOR plasticity mechanisms. Here, we evaluated the mechanisms of induction of MOR-LTD and MOR-STD in the DLS using pharmacology and optogenetics combined with patch clamp electrophysiology. We found that cAMP/PKA signaling and protein synthesis are necessary for MOR-LTD expression, similar to previous studies of cannabinoid-mediated LTD in DLS. MOR-STD does not utilize these same mechanisms. We also demonstrated that cannabinoid-LTD occurs at AIC inputs to DLS. However, while cannabinoid-LTD requires mTOR signaling in DLS, MOR-LTD does not. We characterized the role of presynaptic HCN1 channels in MOR-LTD induction as HCN1 channels expressed in AIC are necessary for MOR-LTD expression in the DLS. These results suggest a mechanism in which MOR activation requires HCN1 to induce MOR-LTD, suggesting a new target for pharmacological modulation of synaptic plasticity, providing new opportunities to develop novel drugs to treat alcohol and opioid use disorders.

**Key Points:**

– Mu opioid receptor-mediated long-term depression at anterior insular cortex inputs to dorsolateral striatum involves presynaptic cAMP/PKA signaling and protein translation, similar to known mechanisms of cannabinoid long-term depression.
– Dorsal striatal cannabinoid long-term depression also occurs at anterior insular cortex inputs to dorsolateral striatum. Dorsal striatal cannabinoid long-term depression requires mTOR signaling, similar to hippocampal cannabinoid long-term depression, but dorsal striatal mu opioid long-term depression does not require mTOR signaling.
– Mu opioid long-term depression requires presynaptic HCN1 channels at anterior insular cortex inputs to dorsolateral striatum.

## Introduction

The opioid system is expressed throughout the brain(Le Merrer *et al*., 2009) and promotes synaptic plasticity in many brain regions(Bao *et al*., 2007; Drake *et al*., 2007; Iremonger & Bains, 2009; Dacher & Nugent, 2011b, a), including the dorsal striatum (DS)(Lovinger, 2010; Atwood *et al*., 2014a; Hawes *et al*., 2017; Munoz *et al*., 2018; Muñoz *et al*., 2020). The DS is the primary input nucleus of the basal ganglia and can be subdivided in two structures: the dorsomedial striatum (DMS), which controls goal-directed learning, and the dorsolateral striatum (DLS), which regulates habit formation(Hilário & Costa, 2008; Lovinger, 2010; O’Tousa & Grahame, 2014; Burton *et al*., 2015; Corbit & Janak, 2016). Drug and alcohol use shift from being outcome-driven to becoming more stimulus-driven and habitual in nature during addiction development(Hopf & Lesscher, 2014; Corbit & Janak, 2016; Everitt & Robbins, 2016; Shen *et al*., 2018). This transition from flexible goal-directed drug use to more habitual use is paralleled by a transition in the neural activity from the DMS to the DLS(Corbit & Janak, 2016; Shen *et al*., 2018; Renteria *et al*., 2020), thereby shifting the balance of action control from DMS to DLS. Previous reports have demonstrated the role of glutamatergic synaptic plasticity in behavior, showing that the disruption of glutamatergic long-term synaptic depression (LTD) at orbitofrontal cortex-DMS synapses prevents habit learning(Gremel *et al*., 2016), and the disruption of glutamatergic LTD in DLS promotes habit learning(Nazzaro *et al*., 2012; DePoy *et al*., 2013). Therefore, changes in the expression of excitatory synaptic plasticity affect specific DS-related behaviors.

Our previous work established that mu opioid receptor (MOR) activation induces long-term synaptic depression (MOR-LTD) or short-term depression (MOR-STD) of glutamate release in both the DLS and DMS(Atwood *et al*., 2014a; Munoz *et al*., 2018; Muñoz *et al*., 2020). We demonstrated that MOR-mediated synaptic inhibition is expressed at multiple DS synapses (cortical, thalamic, amygdala and cholinergic), but only a subset of corticostriatal synapses expressed MOR-LTD(Atwood *et al*., 2014a) that was disrupted by prior *in vivo* ethanol exposure(Munoz *et al*., 2018; Muñoz *et al*., 2020). Like with alcohol, we also found that mice that were exposed to the opioid oxycodone *in vivo* had ablated corticostriatal MOR-LTD when measured in brain slices(Atwood *et al*., 2014a; Munoz *et al*., 2018; Muñoz *et al*., 2020). MOR plasticity at other non-corticostriatal DS synapses was unaffected by alcohol exposure(Munoz *et al*., 2018; Muñoz *et al*., 2020). Of particular interest, in DLS, alcohol and opioid-sensitive MOR-LTD of glutamatergic input is completely restricted to anterior insular cortex (AIC) inputs(Atwood *et al*., 2014a; Munoz *et al*., 2018). Altogether, these data led us to hypothesize that MOR plasticity utilizes different mechanisms at distinct glutamatergic synapses in DS. As we previously demonstrated that MOR-LTD and cannabinoid LTD (CB-LTD) are mutually occlusive in DLS(Atwood *et al*., 2014a), we predicted that these two forms of LTD would utilize similar mechanisms. cAMP/PKA signaling mediates CB-LTD in the nucleus accumbens, amygdala, cerebellum, and hippocampus(Atwood *et al*., 2014b). In addition, local protein translation in the presynaptic region is enhanced by the activation of CB1 receptor, which is important for the induction of inhibitory LTD in the hippocampus(Younts *et al*., 2016) and likely also in the DS(Yin *et al*., 2006). CB-LTD at GABAergic synapses in the hippocampus is also mediated by mammalian target of rampamycin (mTOR) signaling(Younts *et al*., 2016). Therefore, we sought to test whether MOR-LTD in DLS, and specifically at AIC-DLS synapses is mediated by cAMP/PKA, protein translation and mTOR signaling. As cAMP also regulates hyperpolarization-activated cyclic nucleotide-gated (HCN) channels, these channels can presynaptically regulate glutamate release, and the type 1 HCN (HCN1) channel may be localized to corticostriatal neurons, we also explored a potential role for HCN1 in mediating MOR-LTD in DLS(Moosmang *et al*., 1999; Postea & Biel, 2011; He *et al*., 2014; Huang *et al*., 2017)

In the current study, we probed glutamatergic synapses in the DLS using *ex vivo* mouse brain slice electrophysiology. We used broad electrical stimulation in C57BL/6J mice to test all glutamatergic inputs. We used an optogenetics approach to probe specific glutamate synapses in DLS, using Emx1-Ai32 mice to test all cortical inputs, VGluT2-Ai32 mice to test all thalamic inputs, and C57BL/6J mice stereotaxically injected with an AAV-ChR2 vector into the AIC to specifically probe AIC-DLS synapses. In combination with these tools, we pharmacologically and genetically probed the mechanism involved in the expression of MOR-LTD in the DLS. We demonstrate that cAMP/PKA signaling and protein synthesis are necessary for MOR-LTD expression, but not for MOR-STD. We describe a novel role of presynaptic AIC HCN1 channels in the induction of MOR-LTD in DLS. We also report that mTOR signaling is required for CB-LTD of glutamate transmission in the DLS, but this is not required for MOR-LTD indicating a divergence in the mechanisms of these two similar forms of synaptic plasticity.

## Methods

### Ethical Approval

Animal care and experimental protocols for this study were approved by the Institutional Animal Care and Use Committee (IACUC) at the Indiana University School of Medicine and all guidelines for ethical protocols and care of experimental animals established by the NIH (National Institutes of Health, Maryland, USA) were followed.

### Methods details

All experiments were performed similar to our previous dorsal striatal electrophysiological studies with some experiment-specific modifications(Atwood *et al*., 2014a; Fritz *et al*., 2018; Munoz *et al*., 2018; Fritz *et al*., 2019). These methods are described in brief below.

### Animals and materials

Male C57BL/6J mice were obtained from the Jackson Laboratory (JAX #000664, Bar Harbor, Maine, USA). Emx1Cre-Ai32, VGluT2Cre-Ai32, and HCN1-flox mice were bred and genotyped in-house (Original stock strains: Ai32: JAX #012569; Emx1Cre: JAX #005628; VGluT2Cre: JAX #016963; HCN1f/f: JAX #028299). All mutant mice used in these studies were backcrossed to C57BL/6J mice for a minimum of 7 generations. The mice used in these studies were between PND 60-100 at the time of experimentation (with the exception of HCN1-flox AAV-cre-injected mice ~PND 98-154). Animals were group-housed in a standard 12-h light/dark cycle (lights on at 0800) at 50% humidity. Food and water were available *ad libitum*.

### Reagents

We used the MOR agonist [D-Ala^2^, NMe-Phe^4^, Gly-ol^5^]-enkephalin (DAMGO; H-2535, Bachem), GABA_A_ receptor antagonist picrotoxin (PTX, P1675, Sigma-Aldrich), PKI (6221, Tocris), KT5720 (1288, Tocris), Forskolin (F3917, Sigma-Aldrich), Cycloheximide (0970, Tocris), ZD7288 (1000, Tocris), WIN55,212-2 (1038, Tocris), and Torin 2 (4248, Tocris). Other reagents used for making solutions were purchased from Sigma-Aldrich or Fisher Scientific.

### Brain slice preparation

Mice were euthanized via decapitation under deep isoflurane anesthesia, and the brain was quickly excised and placed in an ice-cold cutting solution containing (in mM): 194 sucrose, 30 NaCl, 4.5 KCl, 1 MgCl_2_, 26 NaHCO_3_, 1.2 NaH_2_PO_4_, 10 Glucose saturated with a mixture of 95% O_2_ and 5% CO_2_, and sliced to a thickness of 280 μm on a vibratome (Leica VT1200S, Germany). Slices were transferred to an artificial cerebrospinal fluid (aCSF) solution containing (in mM): 124 NaCl, 4.5 KCl, 1 MgCl_2_, 26 NaHCO_3_, 1.2 NaH_2_PO_4_, 10 Glucose, 2 CaCl_2_ (310-320 mOsm) saturated with 95% O_2_/5% CO_2_ at 30°C for 1 hr before being moved to room temperature. When ready for recording, slices were transferred to a recording chamber continuously perfused with aCSF solution saturated with 95% O_2_/5% CO_2_.

### Electrophysiology recordings

Whole-cell recordings of excitatory postsynaptic currents (EPSCs) in medium spiny neurons (MSNs) were carried out at 29–32 C° and aCSF was continuously perfused at a rate of 1-2 ml/min. Recordings were performed in the voltage clamp configuration using a Multiclamp 700B amplifier and a Digidata 1550B (Molecular Devices, San Jose, CA). Slices were visualized on an Olympus BX51WI microscope (Olympus Corporation of America). MSNs were identified by their size, membrane resistance, and capacitance. Picrotoxin (50 μM) was added to the aCSF for recordings to isolate excitatory transmission. Patch pipettes were prepared from filament-containing borosilicate micropipettes (World Precision Instruments) using a P-1000 micropipette puller (Sutter Instruments, Novato, CA), having a 2.0-3.5 MΩ resistance. The internal solution contained (in mM): 120 CsMeSO_3_, 5 NaCl, 10 TEA-Cl, 10 HEPES, 5 lidocaine bromide, 1.1 EGTA, 0.3 Na-GTP and 4 Mg-ATP (pH 7.2 and 290-310 mOsm). MSNs were voltage clamped at −60 mV for the duration of the recordings. For electrically-evoked recordings, a twisted tungsten bipolar stimulating electrode (PlasticsONE, Roanoke, VA) was placed at the border of the white matter of the external capsule. Electrically-evoked excitatory postsynaptic currents (eEPSCs) were generated by a DS3 Isolated Current Stimulator (Digitimer, Ft. Lauderdale, FL) every 20 s and stimulus intensity was adjusted to produce stable eEPSCs of 200–600 pA in amplitude prior to the initiation of experimental recording. To induce cannabinoid-mediated LTD (CB-LTD), high-frequency stimulation (HFS; 4 pulses of 100_Hz, 10_s inter-pulse interval) coupled with depolarization (0 mV) was delivered after 10 minutes of baseline recording(Atwood *et al*., 2014a; Munoz *et al*., 2018). To induce hippocampal LTP, a concentric bipolar stimulating Tungsten electrode (MicroProbes for Life Science, MD) was placed into stratum radiatum in area CA1 ~500 μm from the recording site. The stimulus intensity for these experiments ranged from 200 pA to 1800 pA due to differences in recording conditions. The internal solution contained (in mM): 120 K-gluconate, 10 NaCl, 5 MgCl2, 2 phosphocreatine, 25 HEPES, 0.2 EGTA, 0.3 Na-GTP and 5 Mg-ATP (pH 7.35 and 290-300 mOsm) and the CA1 pyramidal neurons were held at −90mV. After the baseline was established, LTP was induced by switching the mode to current-clamp and delivering high-frequency stimulation (HFS; 4 pulses of 100□Hz, 1 min inter-pulse interval) as previously reported(Duffy & Nguyen, 2003). The spontaneous EPSC (sEPSC) recordings were made for 2 min and the analysis of frequency (Hz), decay constant (ms), rise constant (ms) and amplitude (pA) were measured to determine the effects of the blockade of striatal HCNs and the deletion of HCN1 in AIC. The decay constant of sEPSCs was fitted as single exponential and both rise and decay-phase were fitted between 10 and 90% of the maximal amplitude. Data were acquired using Clampex 10.3 (Molecular Devices, San Jose, CA). Series resistance was monitored and only cells with a stable series resistance (less than 25 MΩ and that did not change more than 15% during recording) were included for data analysis. Recordings were made 2–7 h after euthanasia.

### Viral injections

C57BL/6J mice were anesthetized with isoflurane and stereotaxically-injected with the adeno-associated viral (AAV) vector, AAV9.hSyn.ChR2(H134R)-eYFP (Addgene #26973) to drive ChR2 expression in AIC neurons. Bilateral injections were made into AIC: A/P: +2.4, M/L: ±2.3, D/V: −2.25 (50 nl/injection, 12.5 nl/min infusion rate).

To produce AIC projection neuron HCN1 knockout mice, HCN1-flox mice were anesthetized with isoflurane and stereotaxically-injected with AAV9.hSyn.HI.eGFP.Cre.WPRE.SV40 (Addgene #105540) or AAV9.hSyn.eGFP.WPRE.bGH as control (Addgene #105539). Bilateral injections were made into AIC at coordinates: A/P: +2.4, M/L: ±2.3, D/V: −2.25 (75 nl/injection, 12.5 nl/min infusion rate). HCN1-flox mice were allowed to recover for at least 4 weeks to allow for adequate ablation of HCN1 channel expression before brain slices were made for electrophysiological recordings. Prior to recording, brain slices were imaged via an Olympus MVX10 microscope (Olympus Corporation of America) to verify GFP or GFP-tagged cre-recombinase expression in injected HCN1-flox mice.

### Optogenetic Recordings

AAV-ChR2 injection in C57BL/6J mice was performed to target ChR2 expression to inputs from AIC to DLS. Optically-evoked EPSCs (oEPSCs) in MSNs from Emx1-Ai32, VGluT2-Ai32 and C57BL/6J mice injected with AAV-ChR2 were produced in brain slices using 470-nm blue light (5-ms exposure time) delivered via field illumination through the microscope objective. Light intensity was adjusted to produce stable oEPSCs of 200–600-pA amplitude prior to experimental recording. oEPSCs were evoked every 30s. AIC-DLS CB-LTD was produced using optical high-frequency stimulation (oHFS) coupled with depolarization (4 blue light pulses of 50□Hz, 10□s inter-pulse interval). Prior to recording, brain slices were imaged via an Olympus MVX10 microscope (Olympus Corporation of America) to verify YFP-tagged ChR2, GFP-tagged cre-recombinase, or GFP expression in injected C57BL/6J mice. Injection site location were determined from matching images to the Reference Allen Mouse Brain Atlas (Figs. 2 and 12). Animals that did not have viral expression in the AIC were excluded from the study

### Quantitative polymerase chain reaction

Anterior insular cortex tissues were taken from HCN1-flox/AAV-cre (+) and HCN1-flox/AAV-GFP (−) mice. RNA was isolated from brain tissue using the RNeasy Plus Universal Mini Kit (Qiagen #73404) according to the manufacturer’s protocol. Total RNA (50 ng/μl) was converted to complementary DNA (cDNA) using the High Capacity cDNA Reverse Transcription kit (Catalog number: 468814, Applied Biosystems Inc. (ABI), Foster City, CA) and amplified using a CFX Connect real-Time PCR detection System (Bio-Rad Laboratories). The TaqMan probe used in the current study was Hcn1 (PrimePCR™ Assay: Hcn1.Mmu, Unique Assay ID:qMmuCED0045731, Catalog #10025636, ThermoFisher). Quantitative PCR was performed using SsoAdvanced Universal Probes Supermix (Catalog #: 172-5280, Bio-Rad Laboratories). The relative amount of each transcript was determined via normalization across all samples to the endogenous control glyceraldehyde 3-phosphate dehydrogenase (GAPDH) (Catalog: Mm.PT.39a.1, PrimeTime Std® qPCR Assay, IDT, IA, USA) to account for variability in the initial concentration and quality of the total RNA and in the conversion efficiency of the reverse transcription reaction as recommended by ABI. In addition, before initiation of the analysis, cDNA was diluted 1:10 and amplified using the respective TaqMan probes to ensure that the amount of cDNA used was in the linear range. RNA samples from each individual animal were run in duplicate.

To quantify the relative expression levels of the different genes for each mouse genotype, we calculated the difference (ΔCt) between the cycle threshold of Hcn1 and the housekeeping gene glyceraldehyde 3-phosphate dehydrogenase (GAPDH). From these data, the ΔΔCt ([ΔCt(Cre (+))–ΔCt(GFP (−))]) was computed and converted to a relative quantitative (RQ) value using the formula 2−ΔΔCt. The results were tabulated as mean□±□SD and compared between genotypes via unpaired Student’s t tests.

### Quantification and statistical analysis

#### Sample size

The target number of samples in each group for electrophysiological experiments was determined from our previously published studies(Atwood *et al*., 2014a; Munoz *et al*., 2018; Muñoz *et al*., 2020).

#### Replication

Sample sizes indicated in figures for electrophysiological experiments represent biological replicates. Due to the inherent biological variability between neurons recorded (biochemical makeup and innervation patterns), neurons were considered the unit of biological replication. 1 neuron was recorded per brain slice and all experiments involved recordings from at least 3 mice.

#### Data analyses

eEPSCs and oEPSCs were analyzed using Clampfit 10.1 (Molecular Devices, San Jose, CA) and sEPSCs with MiniAnalysis 6.0 (Synaptosoft Inc). Data are presented as the mean ± SD in the main text and as mean ± SEM (time courses) or as box plots representing medians and interquartile ranges in the figure legends. The analyses of normally distributed data were performed using two-tailed unpaired or two-tailed paired Student’s t tests following an F test to confirm similar variances. Non-normally distributed data were analyzed using two-tailed Welch’s t test for unpaired data (Fig. 7D). One-way ANOVA following Dunnett’s multiple comparisons test was performed to compare controls from treatments in Figs. 4G, N, U and 9M. Repeated measures One-way ANOVA following Tukey’s multiple comparisons test was performed for Fig. 9C, F, I and L. Data that were analyzed using this test are indicated in the figure legends.

Statistical analyses were performed with Prism 9 (GraphPad, La Jolla, CA). The level of significance was set at *P* < 0.05 for all analyses. Representative traces are the average baseline EPSC (1-10 min), average of the acute DAMGO response (12-20 min) and average post-treatment EPSC of final 10 min of recording. Exclusion of individual data points was determined using a ROUT outlier calculator (Q=1%) included in the Prism 9 software package.

## Results

### MOR activation express glutamatergic LTD in the DLS

We previously showed that the activation of MOR in DLS produces LTD of eEPSCs as well as oEPSCs(Munoz *et al*., 2018). Using three approaches, we confirmed that 5□min application of the MOR agonist DAMGO (0.3□μM) induces MOR-LTD. We found that DAMGO produced a persistent decrease in eEPSC amplitude in DLS MSNs (74 ± 10%; Fig. 1A-D) in brain slices from C57Bl/6J mice. To generally probe corticostriatal synapses, we used 470 nm light delivered through the microscope objective to evoke oEPSCs in brain slices from Emx1-Ai32 mice (Fig. 1E), where the activation of MOR again induced MOR-LTD (75 ± 10%; Fig. 1F-H). Finally, to specifically stimulate the release of glutamate from AIC inputs to the DLS, we infused an AAV vector expressing ChR2 in the AIC (Fig. 1I, Fig. 2), as we previously did(Munoz *et al*., 2018). As shown previously, activation of MORs produced glutamatergic LTD of AIC-DLS transmission (76 ± 15%; Fig. 1J-L). These data replicate our previous work that MORs mediated LTD of AIC-DLS synapses. We previously showed that stimulation-induced cannabinoid-mediated LTD (CB-LTD) and MOR-LTD were mutually occlusive in DLS (Atwood *et al*., 2014a). In order to determine whether we could infer potential MOR-LTD mechanisms from what is known about CB-LTD, we investigated whether CB-LTD also occurred at AIC-DLS synapses. We used the same AAV approach described above (Fig. 3A) and tested if we could produce stimulation-induced LTD (that is likely mediated by endocannabinoid signaling) and also tested responses to the CB1 receptor agonist WIN55,212-2 (1 μM) to chemically induce CB-LTD. We found that stimulation-induced LTD (65 ± 7%; Fig. 3B, C, D and F) and chemically-induced CB-LTD (68 ± 7%; Fig. 3B, C, E and F) both occur at AIC-DLS synapses. Next, we explored the signaling cascades that mediate MOR plasticity in DLS using what is known about CB-LTD as a guide for our experiments.

**Figure 1.**
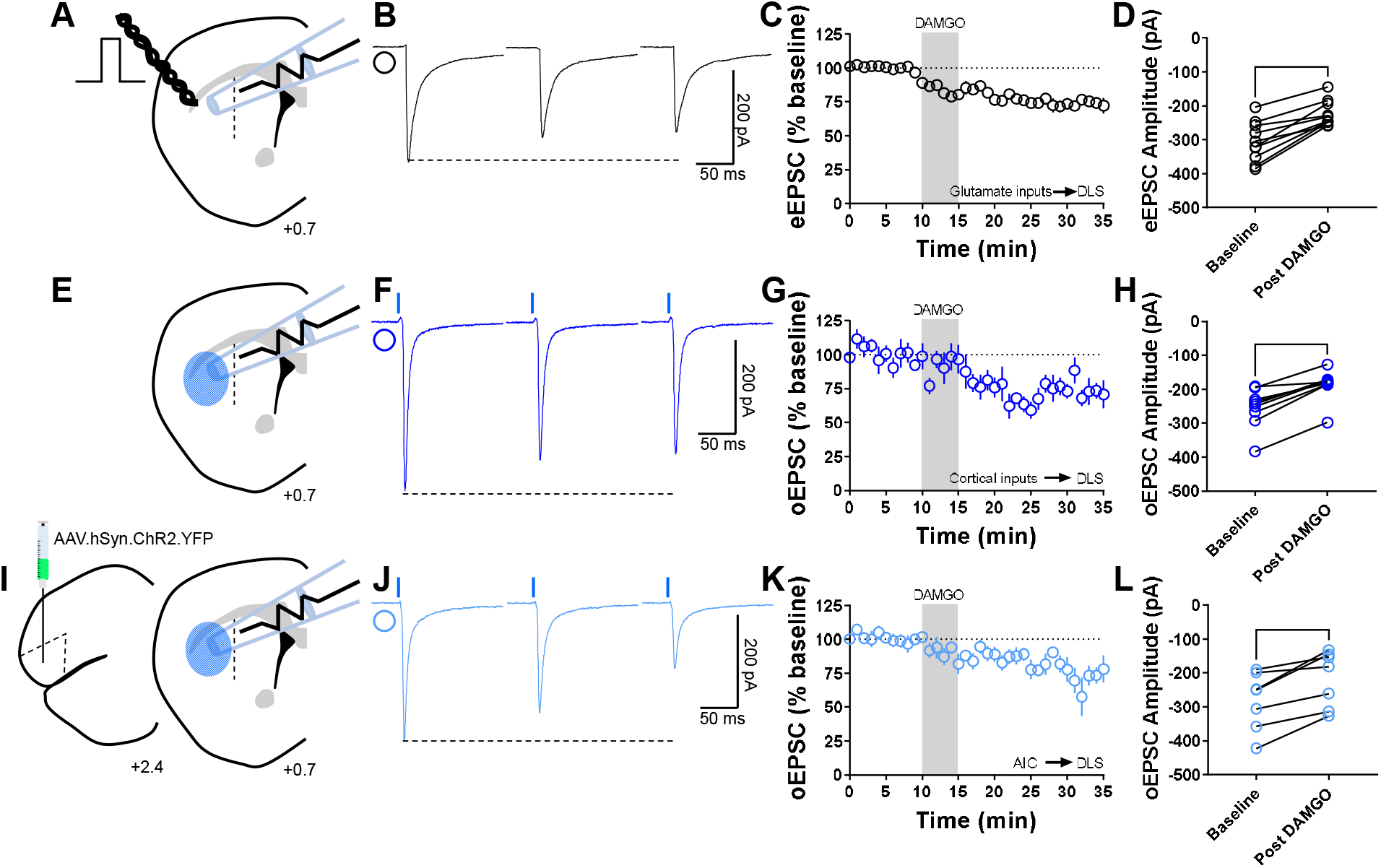
MOR activation induces glutamatergic LTD in the DLS. **(A)** Schematic representation of coronal brain slice showing the recording of EPSCs evoked by focal electric stimulation in the DLS of C57BL/6J mice. **(B)** Representative electrically-evoked EPSC traces before, during and after DAMGO (0.3□μM, 5□min) application. **(C)** The activation of MOR by DAMGO induced glutamatergic LTD in DLS MSNs of C57BL/6J mice (final 10□min of recording average: 74 ± 3%). **(D)** eEPSC amplitudes in MSNs within DLS were significantly reduced after DAMGO application (0–10□min baseline v. final 10□min of recording; paired t-test, P<0.0001, t9□=6.688, n□=□10 neurons from 5 mice). **(E)** Schematic representation of coronal brain slice showing the recording of EPSCs evoked by focal optical stimulation (470-nm blue light for 5-ms exposure) in the DLS of Emx1-Ai32 mice. **(F)** Representative optically-evoked EPSC traces before, during and after DAMGO (0.3□μM, 5□min) application. **(G)** The activation of MOR by DAMGO induced corticostriatal LTD in DLS MSNs of Emx1-Ai32 (final 10□min of recording average: 75 ± 3%). **(H)** oEPSC amplitudes were significantly reduced after DAMGO application (0–10□min baseline v. final 10□min of recording; paired t-test, P<0.0001, t8□=7.73, n□=□9 neurons from 4 mice). **(I)** Schematic figure of the injection paradigm showing an AAV vector encoding for ChR2 (AAV.hSyn.ChR2.YFP) in AIC in C57BL/6J mice, this AAV was injected 2 weeks prior to recordings. Also, the next schematic representation of coronal brain slice shows the recording of oEPSCs (470-nm blue light for 5-ms exposure) in the DLS. **(J)** Representative AIC-DLS oEPSC traces before, during and after DAMGO (0.3□μM, 5□min) application. **(K)** DAMGO induced AIC-DLS LTD (final 10□min of recording average: 76 ± 5%). **(L)** DAMGO application significantly reduced oEPSC amplitudes (0–10□min baseline v. final 10□min of recording; paired t-test, P□=□0.00403, t6□=4.5, n□=□7 neurons from 4 mice). Data represent mean ± SEM. **P□<□0.01, ****P□<□0.0001.

**Figure 2.**
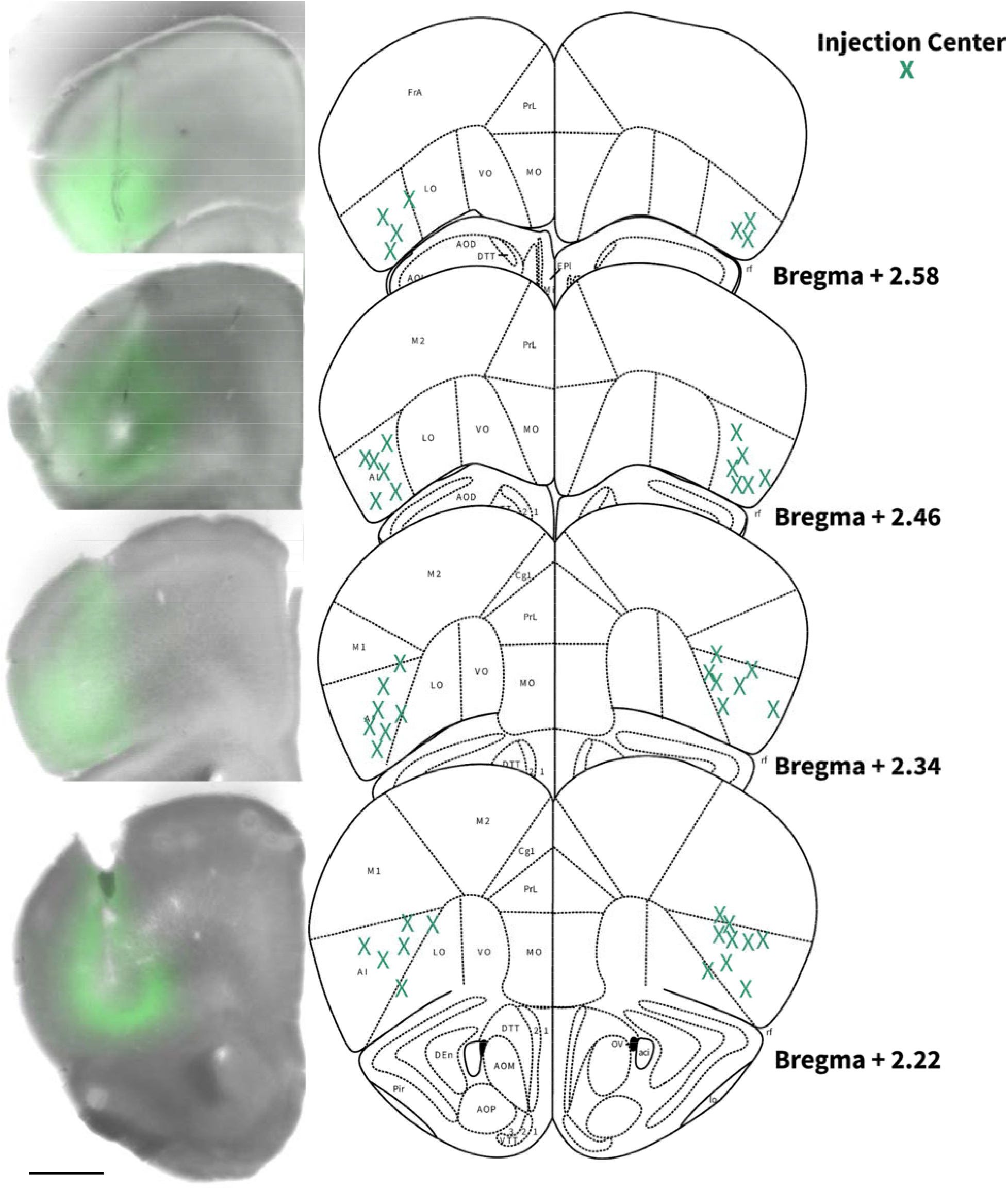
Locations of AIC injections for DLS electrophysiology. Coronal brain hemi slices images and schematic figures modified from Allen Mouse Brain Atlas showing bilateral injection centers of AAV-ChR2-EYFP in the AIC of C57BL/6J mice verified by histology.

**Figure 3.**
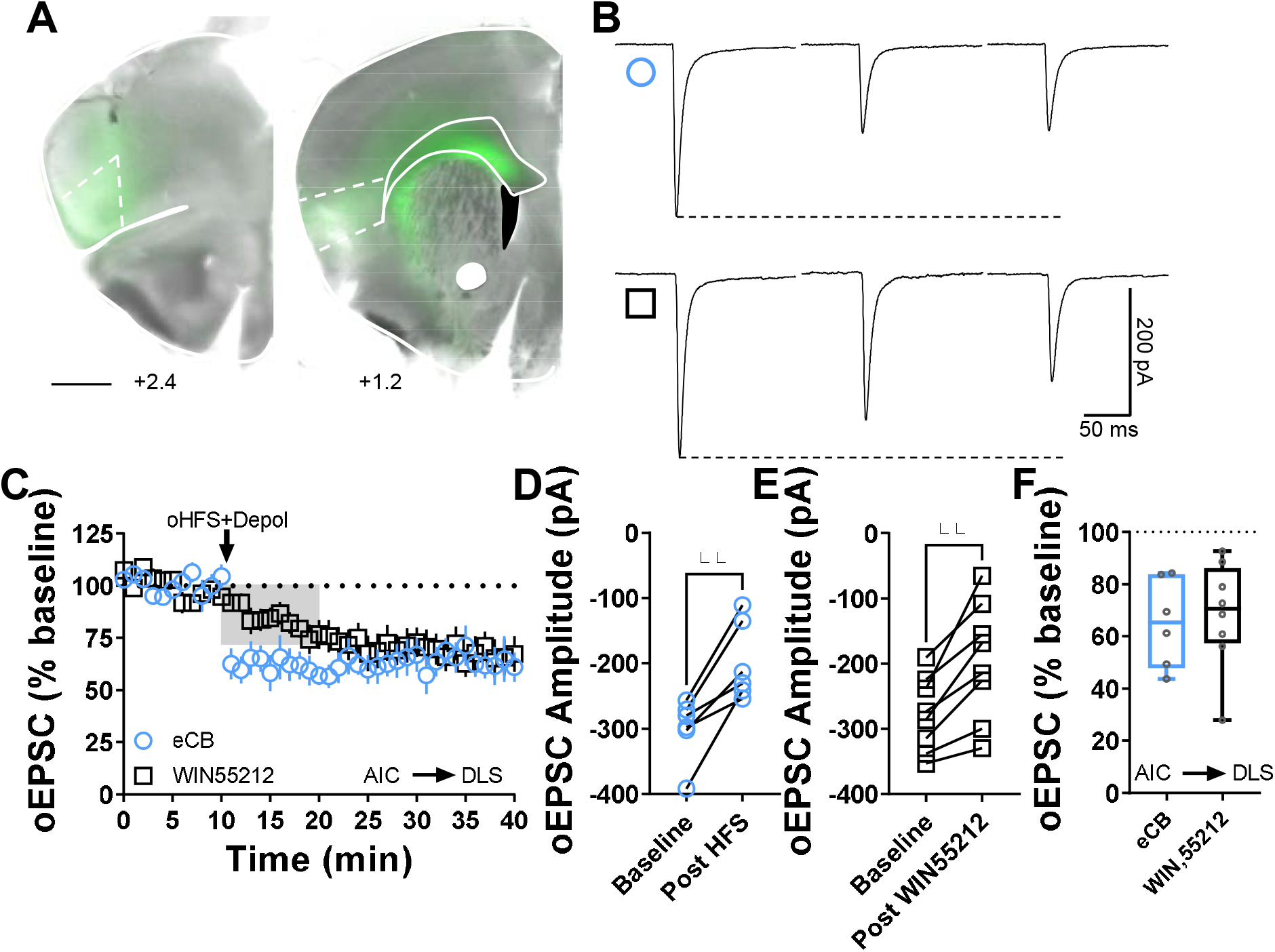
Cannabinoid LTD occurs at AIC-DLS synapses. **(A)** Coronal brain slice showing the AAV-ChR2 infection of AIC and dorsal striatal terminal expression (bar scale□=□1000□μm). **(B)** Representative oEPSC traces showing the effects of pairing optical high frequency stimulation with postsynaptic depolarization (oHFS+Depol, cyan open circle) to physiologically induce LTD, presumably mediated by endocannabinoids, and the application of the CB1 agonist, WIN55,212-2 (1□μM, 10□min, black open square) to chemically induce CB1-LTD in brain slices of AAV-ChR2 C57Bl/6J mice. **(C)** Both oHFS and WIN55,212-2 induced AIC-DLS LTD. Data represent mean ± SEM. **(D)** oHFS significantly reduced oEPSC amplitudes (0–10□min baseline v. final 10□min of recording; paired t-test, P□=□0.00350, t5=5.2, n□=□6 neurons from 4 mice). **(E)** WIN55,212-2 significantly reduced oEPSC amplitudes (0–10□min baseline v. final 10□min of recording; paired t-test, P□=□0.00179, t7=4.9, n□=□8 neurons from 4 mice). **(F)** The box plot shows the final 10□min of recording average of stimulation-induced LTD (65 ± 7%) and CB1-LTD (68 ± 7%). **P□<□0.01.

### Inhibition of presynaptic PKA blocks MOR-mediated LTD

The activation of opioid receptors produces an inhibition of adenylyl cyclase (AC) mediated by Gαi, decreasing cAMP levels, which leads to an attenuation of cAMP/protein kinase A (PKA) signaling(Atwood *et al*., 2014b; Reeves *et al*., 2022). However, little is known about PKA’s role in MOR-mediated LTD. Using KT5720 (1 μM), a specific and cell permeable inhibitor of PKA, which brain slices were incubated in for 1h prior to recordings, we demonstrated that the inhibition of PKA blocks the maintenance of MOR-LTD of eEPSCs in DLS (101 ± 19%; Fig. 4A-C and G). Corticostriatal MOR-LTD was also blocked by PKA inhibition (94 ± 16%; Fig. 4H-J and N), as was MOR-LTD at AIC-DLS synapses (94 ± 11%; Fig. 4O-Q and U). We found that these effects were due to inhibiting presynaptic PKA activity as at least 30 min treatment with the membrane-impermeable PKA inhibitor PKI (1 μM), intracellularly infused through the recording micropipette into the postsynaptic MSN, did not affect MOR-LTD in the DLS (eEPSCs: 84 ± 7%; Fig. 4D-G) (corticostriatal oEPSCs: 64 ± 20%; Fig. 4K-N) (AIC-DLS oEPSCs: 69 ± 11%; Fig. 4R-U). Since this was a negative effect, we needed to test if PKI was able to induce any effect, therefore, we evaluated PKI in a brain region where the inhibition of postsynaptic PKA disrupts synaptic plasticity(Duffy & Nguyen, 2003). We found that intracellular PKI application blocked the expression of long-term potentiation (LTP) in hippocampal CA1 pyramidal neurons (Control: 205 ± 49% vs PKI: 88 ± 9%, Fig. 5). As we expected, our data demonstrate the role of presynaptic PKA in MOR-LTD.

**Figure 4.**
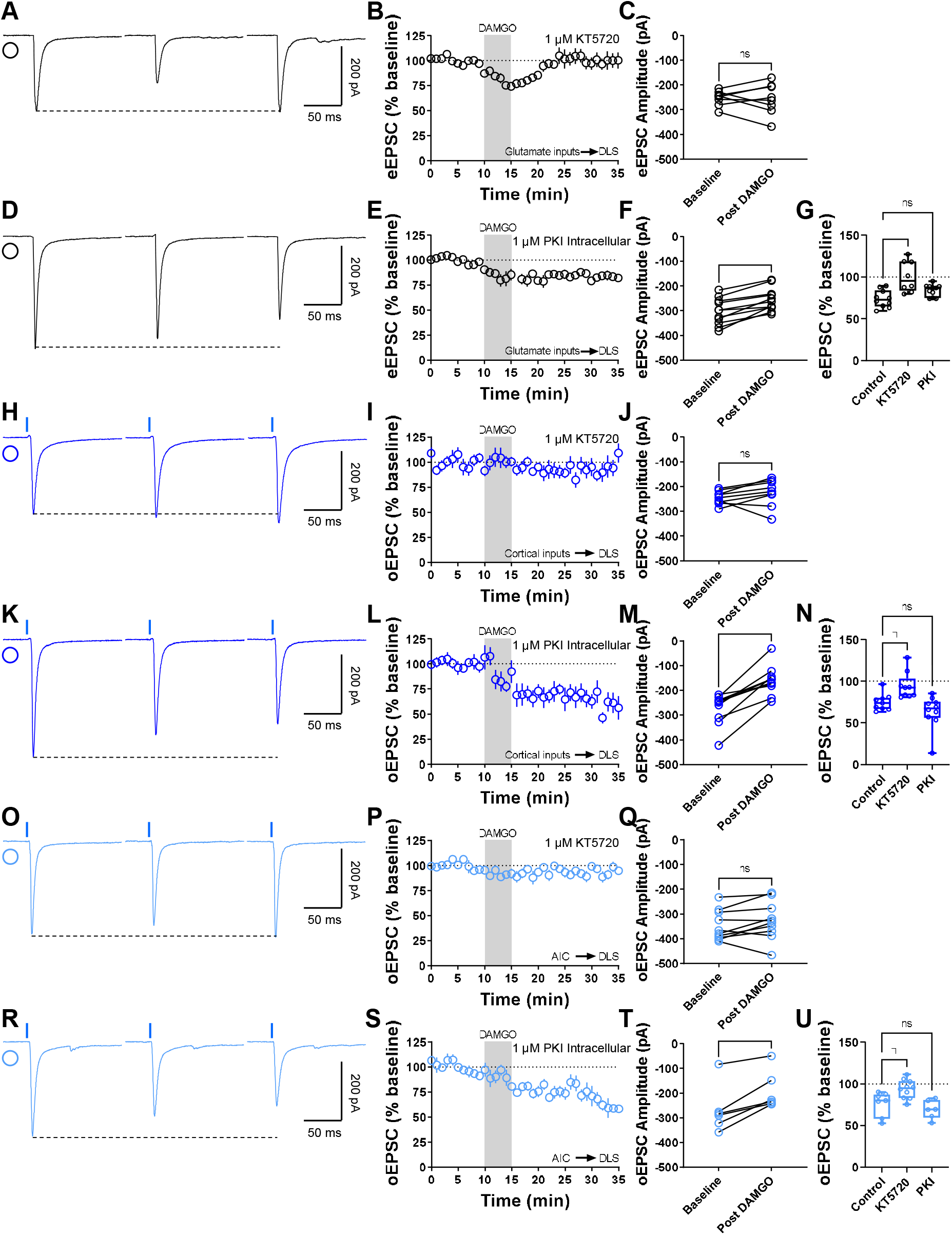
Inhibition of presynaptic PKA blocks MOR-mediated LTD. **(A)** Representative eEPSC traces showing the effects of DAMGO (0.3□μM, 5□min) application after the preincubation of PKA-selective inhibitor, KT5720 (1 μM, ≥1 hr). **(B-C)** The preincubation of KT5720 blocked glutamatergic MOR-LTD, the eEPSC amplitudes did not change after DAMGO application (0–10□min baseline v. final 10□min of recording; paired t-test, P□=□0.865, t7□=0.176, n□= 8 neurons from 3 mice). **(D)** Representative eEPSC traces showing the effects of DAMGO (0.3□μM, 5□min) application after the intracellular dialysis of PKI (1 μM, ≥30 min). **(E-F)** The inhibition of postsynaptic PKA did not alter MOR-LTD. eEPSC amplitudes were reduced after DAMGO application (0–10□min baseline v. final 10□min of recording; paired t-test, P□=□0.000120, t10□=6.07, n□= 11 neurons from 5 mice). **(G)** Presynaptic PKA inhibition disrupted MOR-LTD (KT5720: 101 ± 7%, P=0.000193 v PKI: 84 ± 2%, P=0.121; F(2,26)=10.65, One-way ANOVA Dunnet’s multiple comparison test) **(H)** Representative oEPSC traces showing the effects of DAMGO (0.3□μM, 5□min) application after the preincubation of PKA-selective inhibitor, KT5720 (1 μM, ≥1 hr). **(I-J)** PKA inhibition blocked corticostriatal MOR-LTD, with no changes in oEPSC amplitudes after DAMGO application (0–10□min baseline v. final 10□min of recording; paired t-test, P□=□0.183, t8□=1.46, n□= 9 neurons from 4 mice). **(K)** Representative oEPSC traces showing the effects of DAMGO (0.3□μM, 5□min) application after the intracellular dialysis of PKI (1 μM, ≥30 min). **(L-M)** The inhibition of postsynaptic PKA did not alter MOR-LTD. oEPSC amplitudes were reduced after DAMGO application (0–10□min baseline v. final 10□min of recording; paired t-test, P□=□0.000118, t9□=6.45, n□= 10 neurons from 4 mice). **(N)** Presynaptic PKA inhibition disrupted cortical MOR-LTD (KT5720: 94 ± 5%, P=0.0305 v PKI: 64 ± 6%, P=0.235; F(2,25)=8.805, One-way ANOVA Dunnet’s multiple comparison test). **(O)** Representative AIC-DLS oEPSC traces showing the effects of DAMGO (0.3□μM, 5□min) application after the preincubation of PKA-selective inhibitor, KT5720 (1 μM, ≥1 hr). **(P-Q)** KT5720 blocked AIC-expressed MOR-LTD, with no changes in oEPSC amplitudes after DAMGO application (0–10□min baseline v. final 10□min of recording; paired t-test, P□=□0.141, t9□=1.613, n□= 10 neurons from 3 mice). **(R)** Representative AIC-DLS oEPSC traces showing the effects of DAMGO (0.3□μM, 5□min) application after the intracellular dialysis of PKI (1 μM, ≥30 min). **(S-T)** Postsynaptic PKA inhibition did not alter specific AIC MOR-mediated LTD. oEPSC amplitudes were reduced after DAMGO application (0–10□min baseline v. final 10□min of recording; paired t-test, P□=□0.00488, t5□=4.8, n□= 6 neurons from 3 mice). **(U)** Presynaptic PKA inhibition disrupted AIC MOR-LTD (KT5720: 94 ± 4%, P=0.0171 v PKI: 69 ± 5%, P=0.518; F(2,20)=8.411, One-way ANOVA Dunnet’s multiple comparison test). Time course data represent mean ± SEM. Box plots show average of the final 10 minutes of recording and represent median and interquartile ranges. ns= not significant, **P□<□0.01, ***P□<□0.001.

**Figure 5.**
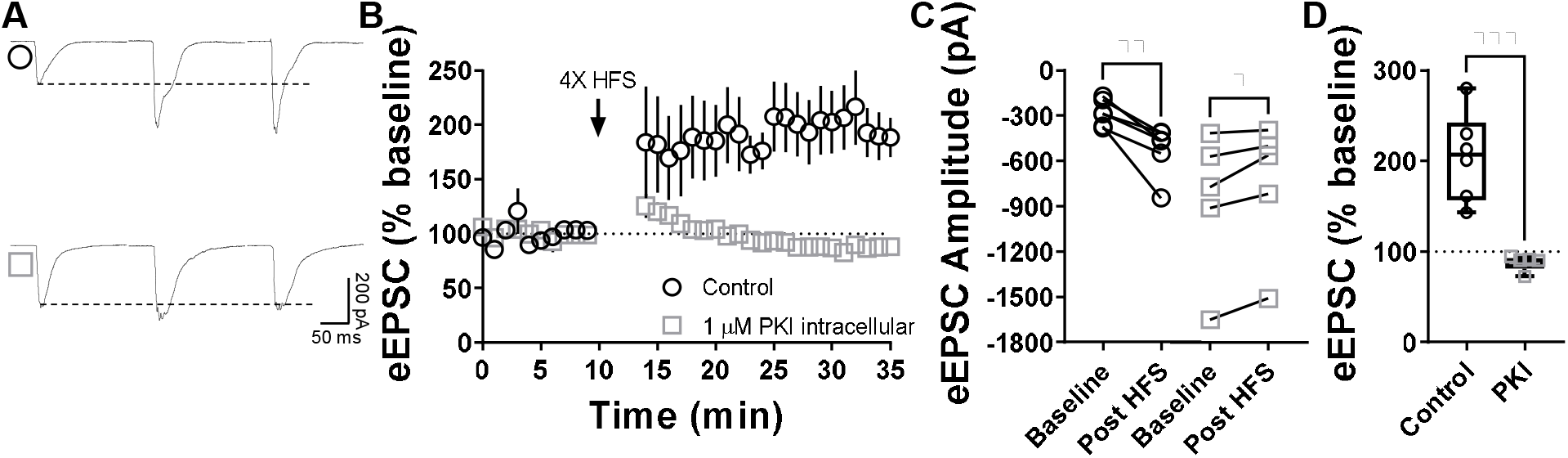
Postsynaptic PKA is required for CA1 hippocampal LTP. **(A)** Representative eEPSC traces before and after high-frequency stimulation (HFS) (4 pulses of 100□Hz, 1□min inter-pulse interval), showing the effects of PKI (1 μM) intracellular diffusion into CA1 pyramidal neurons within hippocampal brain slices of C57BL/6J mice. **(B-C)** Post synaptic PKA signaling is necessary to the induction of LTP in CA1 (0–10□min baseline v. final 10□min of recording; Control: paired t-test, P□=□0.00587, t5□=4.6, n□= 6 neurons from 4 mice; PKI: paired t-test, P□=□0.0282, t4□=3.36, n□= 5 neurons from 4 mice). **(D)** Box plot shows the average disruption of hippocampal LTP by inhibiting post synaptic PKA (Control: 205 ± 49 % vs PKI: 88 ± 9%, P=0.000551, t9=5.22, Unpaired t-test). Time course data represent mean ± SEM. Box plots represent median and interquartile ranges. *P□<□0.05, **P□<□0.01, ***P□<□0.001.

### The activation of adenylyl cyclase disrupts MOR-mediated LTD

Demonstrating a role for PKA signaling in MOR-LTD implicates adenylyl cyclase (AC) signaling as mediating MOR-LTD. To more specifically test this, we persistently activated AC using 20 μM forskolin throughout recording. In the presence of forskolin, DAMGO produced a transient increase of the EPSC amplitudes (Fig. 6A, B, E, F, I and J), followed by a blockade MOR-LTD at all excitatory synapses (105 ± 14%; Fig. 6A-D) and at corticostriatal (105 ± 6%; Fig. 6E-H) and AIC inputs (98 ± 14%; Fig. 6I-L). Altogether, these findings indicate that cAMP/PKA signaling pathway is involved in the induction and maintenance of MOR-LTD in DLS.

**Figure 6.**
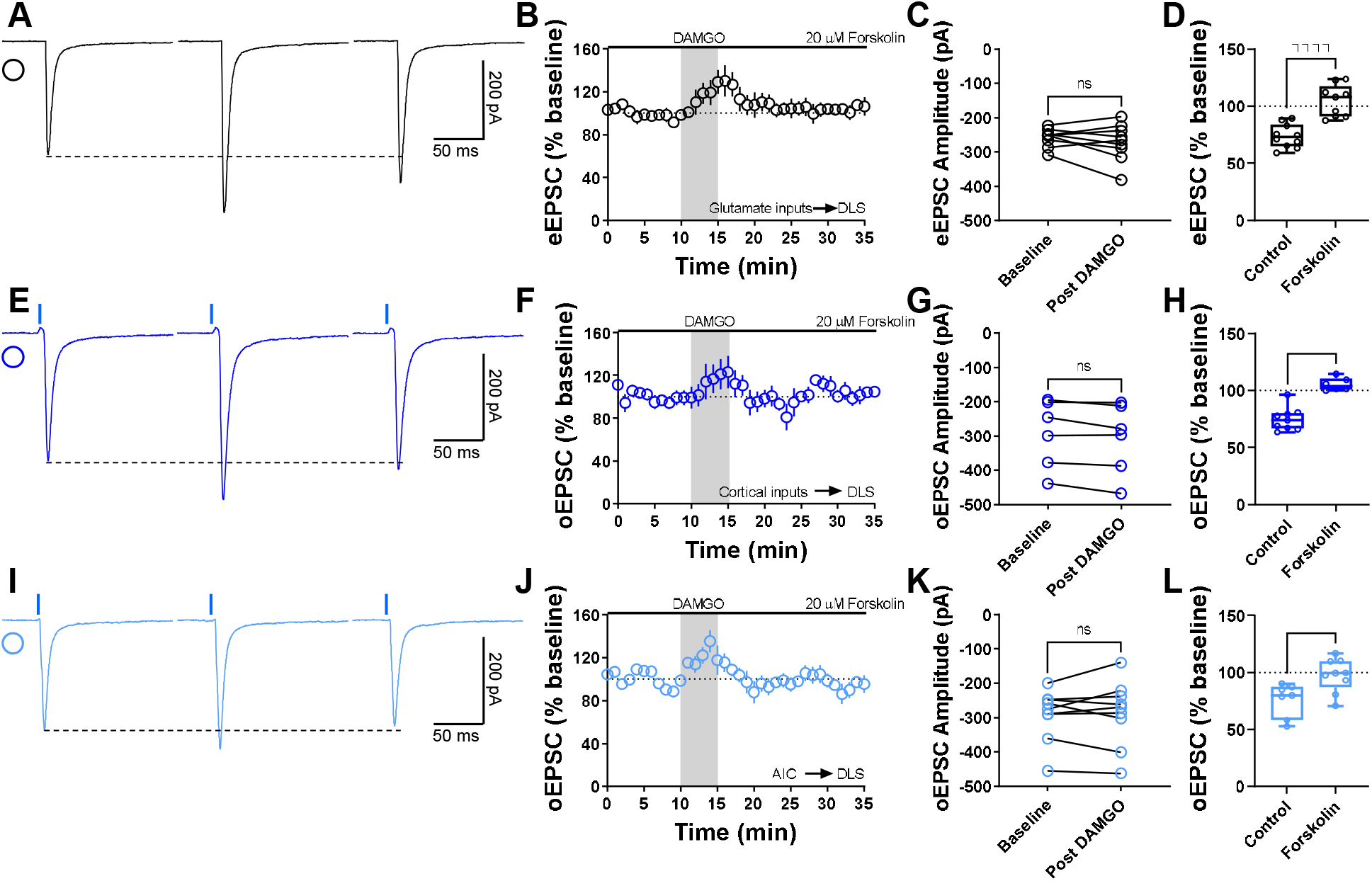
The activation of adenylyl cyclase disrupts MOR-mediated LTD. **(A)** Representative eEPSC traces showing the effects of 20 μM forskolin before, during and after DAMGO (0.3□μM, 5□min) application. **(B-D)** AC activation disrupted glutamatergic MOR-LTD (105 ± 5%, Unpaired t-test, P<0.0001, t17=5.54), showing no changes in eEPSC amplitude after DAMGO bath application (0–10□min baseline v. final 10□min of recording; paired t-test, P□=□0.324, t8□=1.05, n□= 9 neurons from 4 mice). **(E)** Representative oEPSC traces showing the effects of 20 μM forskolin before, during and after DAMGO (0.3□μM, 5□min) application in brain slices from Emx1-Ai32 mice. **(F-H)** AC activation disrupted corticostriatal MOR-LTD (105 ± 2%, Unpaired t-test, P<0.0001, t13=6.65), with no effects in oEPSC amplitude after DAMGO bath application (0– 10□min baseline v. final 10□min of recording; paired t-test, P□=□0.0576, t5□=2.455, n□= 6 neurons from 4 mice). **(I)** Representative AIC-DLS oEPSC traces showing the effects of 20 μM forskolin before, during and after DAMGO (0.3□μM, 5□min) application. **(J-L)** AC activation blocked specifically AIC MOR-LTD (98 ± 5%, Unpaired t-test, P=0.0116, t14=2.9), without changes in oEPSC amplitude after DAMGO bath application (0–10□min baseline v. final 10□min of recording; paired t-test, P□=□0.702, t8□=0.396, n□= 9 neurons from 3 mice). Time course data represent mean ± SEM. Box plots show average of the final 10 minutes of recording and represent median and interquartile ranges. ns= not significant, *P□<□0.05, ****P□<□0.0001.

### Protein translation is required to produce MOR-mediated LTD

Next, we tested the role of protein translation in MOR-LTD. Using, bath application of 80 μM cycloheximide (selective inhibitor of protein synthesis) during the entire recording, we found that the inhibition of protein translation blocked MOR-LTD generally (105 ± 33%; Fig. 7A-D) and specifically at cortical (102 ± 18%; Fig. 7E-H) and AIC inputs (94 ± 10%; Fig. 7I-L).

**Figure 7.**
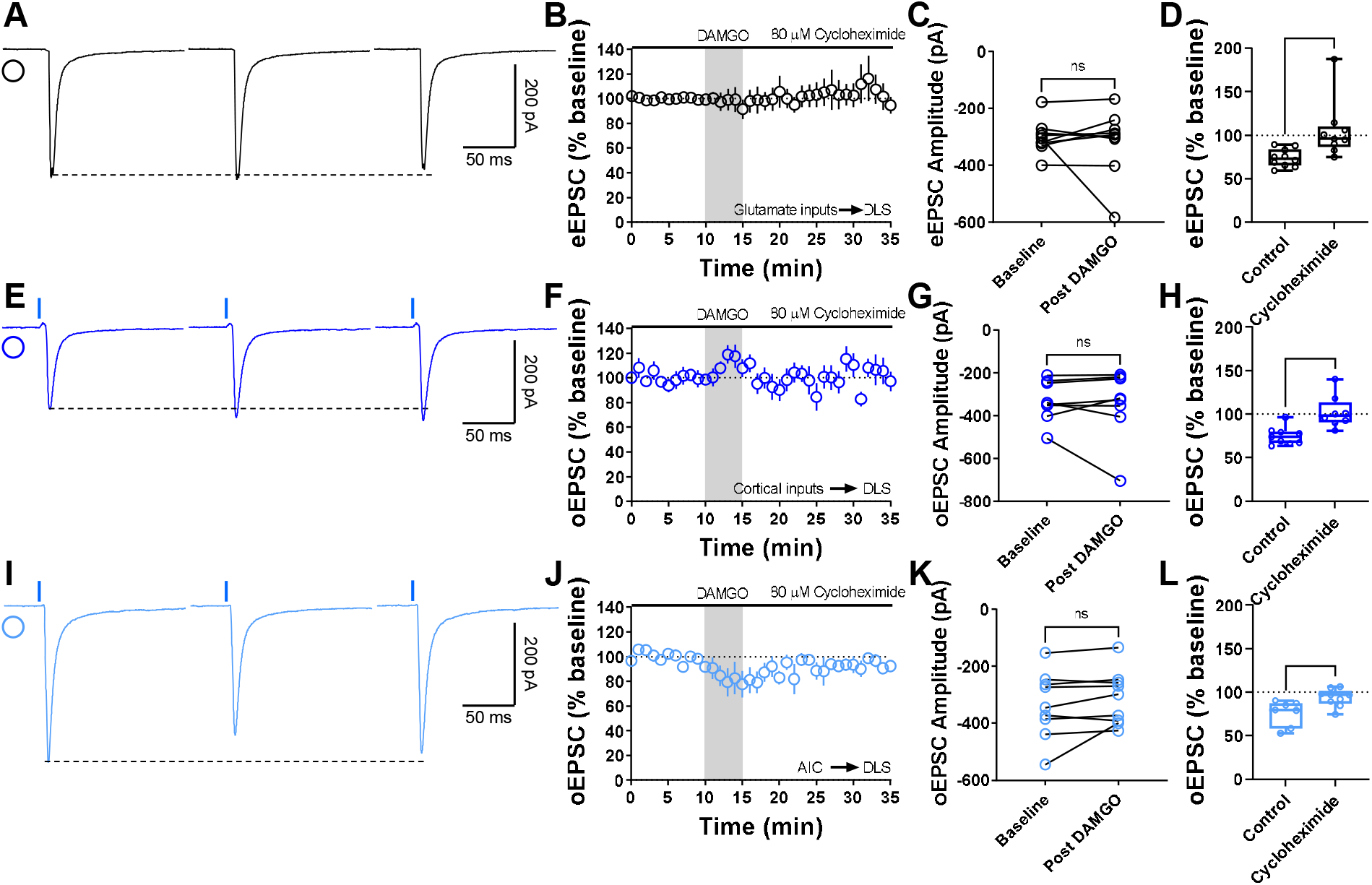
Protein translation is required for MOR-mediated LTD expression. **(A)** Representative eEPSC traces showing the effects of 80 μM cycloheximide before, during and after DAMGO (0.3□μM, 5□min) application. **(B-D)** Protein translation inhibition disrupted glutamatergic MOR-LTD (105 ± 11%, Unpaired Welch’s t-test, P=0.0228, t10=2.72), showing no changes in eEPSC amplitude after DAMGO bath application (0–10□min baseline v. final 10□min of recording; paired t-test, P□=□0.651, t8□=0.471, n□= 9 neurons from 4 mice). **(E)** Representative oEPSC traces showing the effects of 80 μM cycloheximide before, during and after DAMGO (0.3□μM, 5□min) application in brain slices from Emx1-Ai32 mice. **(F-H)** Protein translation inhibition blocked corticostriatal MOR-LTD (102 ± 7%, Unpaired t-test, P=0.00163, t15=3.83) with no effects in oEPSC amplitude after DAMGO bath application (0–10□min baseline v. final 10□min of recording; paired t-test, P□=□0.633, t7□=0.499, n□= 8 neurons from 3 mice). **(I)** Representative AIC-DLS oEPSC traces showing the effects of 80 μM cycloheximide before, during and after DAMGO (0.3□μM, 5□min) application. **(J-L)** Protein translation is needed to induce MOR-LTD (94 ± 3%, Unpaired t-test, P=0.0129, t14=2.85) from AIC inputs. DAMGO did not change in oEPSC amplitude (0–10□min baseline v. final 10□min of recording; paired t-test, P□=□0.163, t8□=1.54, n□= 9 neurons from 3 mice). Time course data represent mean ± SEM. Box plots show average of the final 10 minutes of recording and represent median and interquartile ranges. ns= not significant, *P□<□0.05, **P□<□0.01.

It had been reported that protein translation-mediated CB-LTD involves mTOR signaling in the hippocampus(Younts *et al*., 2016). To test if mTOR is also required for CB-LTD in the DLS, we used bath application of 100 nM Torin 2 (a selective mTOR inhibitor). We demonstrated that the inhibition of mTOR signaling disrupted CB-LTD in the DLS (Control: 76 ± 8% vs Torin 2: 95 ± 5%; Fig. 8A-D). However, MOR-LTD does not require mTOR signaling in the DLS (79 ± 7%; Fig. 8E-H). These data suggest that protein translation is necessary for MOR-mediated LTD, but mTOR signaling is not involved and indicate that CB- and MOR-LTD, while qualitatively similar, do not utilize identical signaling mechanisms.

**Figure 8.**
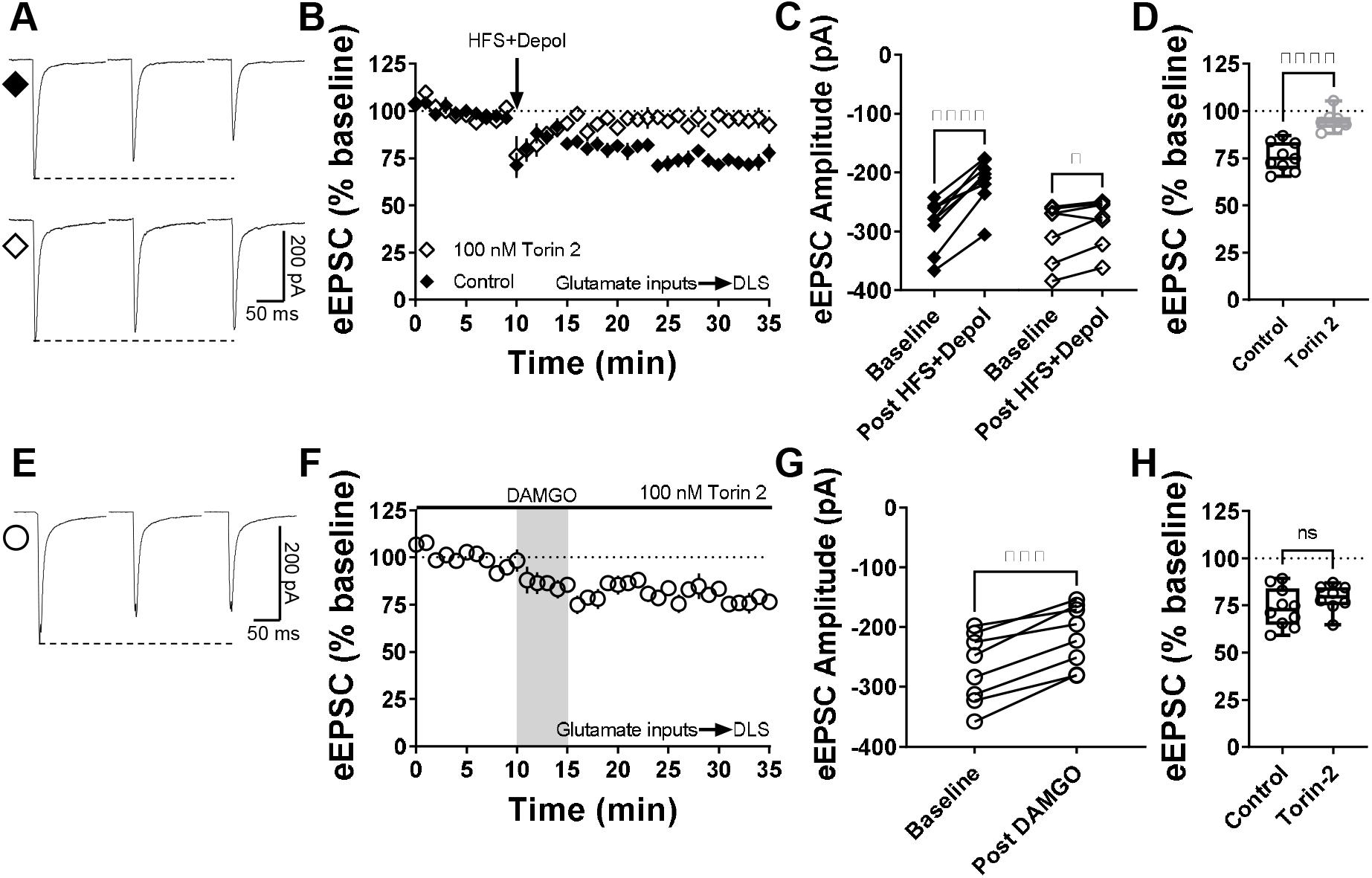
mTOR pathway is involved in cannabinoid-LTD but not in MOR-LTD in the DLS. **(A)** Representative eEPSC traces before and after high-frequency stimulation (HFS) coupled with depolarization (4 pulses of 100□Hz, 10□s inter-pulse interval), showing the effects of torin 2 (100 nM) bath application in brain slices of C57BL/6J mice. **(B-C)** mTOR inhibition blocked this stimulation-induced cannabinoid-LTD in DLS (0–10□min baseline v. final 10□min of recording; Control: paired t-test, P<0.0001, t8□=8.6, n□= 9 neurons from 4 mice; torin 2: paired t-test, P□=□0.0396, t6□=2.62, n□= 7 neurons from 3 mice). **(D)** Torin 2 disrupted stimulation-induced LTD in the DLS (control: 76 ± 3% vs torin 2: 95 ± 2, Unpaired t-test, P<0.0001, t14=5.64) **(E)** Representative eEPSC traces before and after DAMGO bath application (0.3□μM, 5□min), showing the effects of torin 2 (100 nM) bath application in brain slices of C57BL/J. **(F-G)** mTOR inhibition did not affect the expression of MOR-LTD in DLS (0– 10□min baseline v. final 10□min of recording; paired t-test, P□=□0.000140, t7□=7.48, n□= 8 neurons from 3 mice). **(H)** Torin 2 did not blocked MOR-LTD in the DLS (control: 74 ± 3% vs torin 2: 79 ± 3, Unpaired t-test, P=0.270, t16=1.19). Time course data represent mean ± SEM. Box plots show average of the final 10 minutes of recording and represent median and interquartile ranges. ns = not significant, *P<0.05, ***P□<□0.001, ****P□<□0.0001.

### MOR-STD from thalamocortical inputs is not affected by PKA inhibition, cAMP activation or protein translation inhibition

We also confirmed that thalamostriatal inputs expressed MOR-STD in the DLS using VGluT2-Ai32 transgenic mice (12-20 min: 68 ± 5%; 30-35 min: 93 ± 7%; Fig. 9A-B and M) as we previously showed(Atwood *et al*., 2014a; Munoz *et al*., 2018). These data replicate our previous work that MORs mediated STD of thalamostriatal synapses in the DLS. Unlike corticostriatal inputs, these thalamostriatal inputs, were not affected by broad inhibition of PKA (12-20 min: 81 ± 15%; 30-35 min: 108 ± 26%). Inhibition of postsynaptic PKA had no effect on MOR-STD (12-20 min: 86 ± 9%; 30-35 min: 115 ± 31%; Fig. 9D-F and M), just as it had no effect on MOR-LTD. MOR-mediated STD at thalamic synapses was also not affected by the application of forskolin (12-20 min: 81 ± 13%; 30-35 min: 89 ± 15%; Fig. 9G-I and M) or cycloheximide treatment (12-20 min: 76 ± 18%; 30-35 min: 101 ± 23%; Fig. 9J-M). Altogether, these data indicate that MOR-STD at thalamic inputs to DLS does not utilize the same mechanisms that MOR-LTD does.

**Figure 9.**
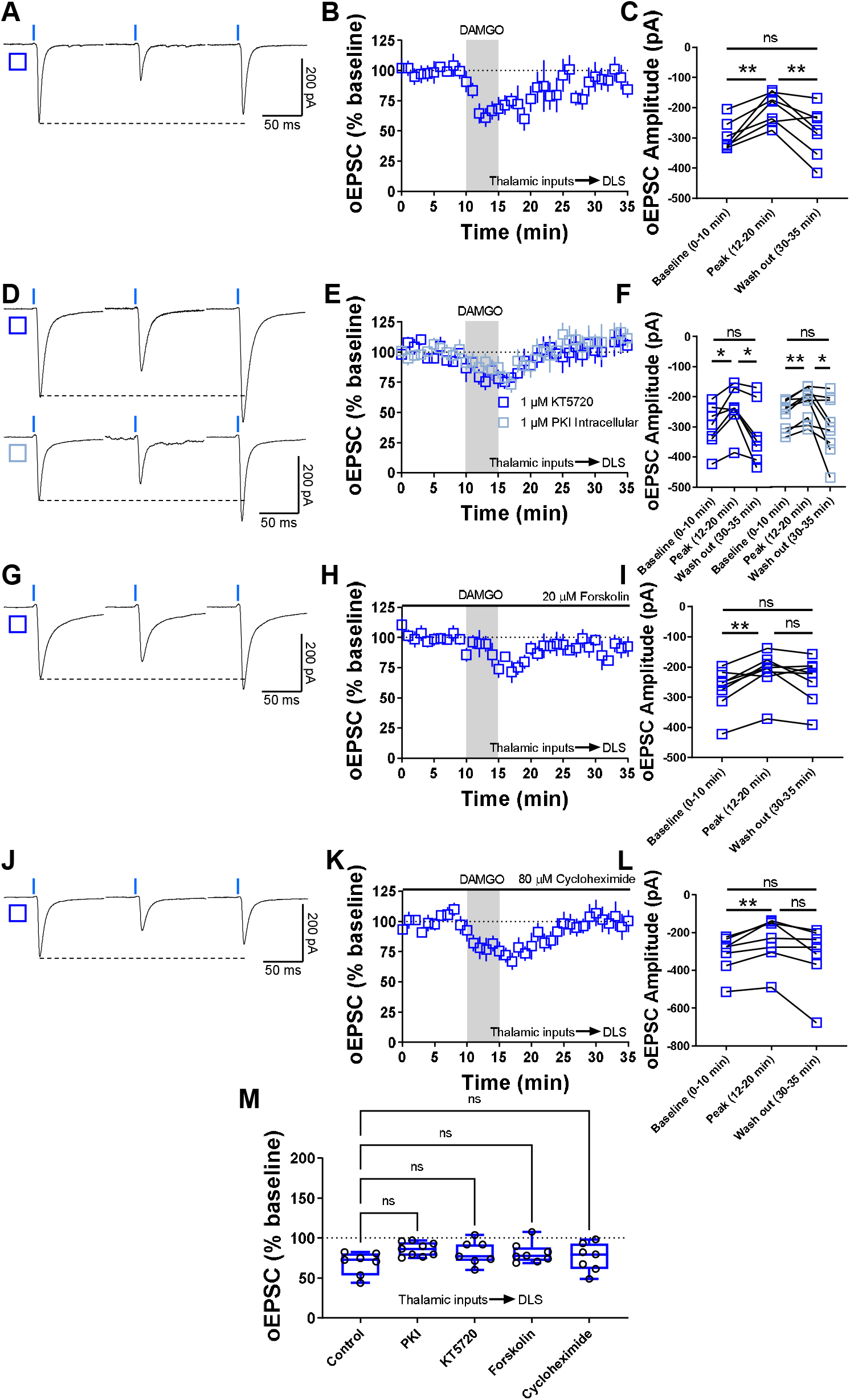
MOR-STD is not affected by PKA inhibition, cAMP activation or protein translation inhibition. **(A)** Representative optically-evoked EPSC traces before, during and after DAMGO (0.3□μM, 5□min) application in brain slices from VGluT2-Ai32 mice. **(B-C)** Thalamocortical inputs produced MOR-STD in DLS from VGluT2-Ai32 mice (10□min baseline v. peak 12-20□min of recording P=0.00275; 12-20□min Peak v 30-35 min Wash out of recording, P=0.0889, Repeated measure One-way ANOVA Tukey’s test, n□=□7 neurons from 3 mice). **(D)** Representative oEPSC traces showing the effects of DAMGO (0.3□μM, 5□min) application after the preincubation of PKA-selective inhibitor KT5720 (1 μM, ≥1 hr, blue open square) and after the intracellular dialysis of PKI (1 μM, ≥30 min, cyan open square). **(E-F)** The inhibition of pre (1 μM KT5720) and postsynaptic PKA (1 μM PKI) did not affected MOR-STD (KT5720: 10□min baseline v. peak 12-20□min of recording, P=0.0353; 12-20□min Peak v 30-35 min Wash out of recording, P□=□0.0353, Repeated measure One-way ANOVA Tukey’s test, n□=□7 neurons from 3 mice) (PKI:, 10□min baseline v. peak 12-20□min of recording, P□=□0.00218; 12-20□min Peak v 30-35 min Wash out of recording, P=0.0429, Repeated measure One-way ANOVA Tukey’s test, n□=□9 neurons from 3 mice). **(G)** Representative oEPSC traces showing the effects of 20 μM forskolin before, during and after DAMGO (0.3□μM, 5□min) application in brain slices from VGluT2-Ai32 mice. **(H-I)** MOR-STD was unaffected after the activation of AC by 20 μM forskolin (10□min baseline v. peak 12-20□min of recording, P=0.00260;12-20□min Peak v 30-35 min Wash out of recording, P=0.168, Repeated measure One-way ANOVA Tukey’s test, n□=□8 neurons from 4 mice). **(J)** Representative oEPSC traces showing the effects of 80 μM cycloheximide before, during and after DAMGO (0.3□μM, 5□min) application in brain slices from VGluT2-Ai32 mice. **(K-L)** Protein translation is not necessary for the expression of MOR-STD (10□min baseline v. peak 12-20□min of recording, P=0.00973;12-20□min Peak v 30-35 min Wash out of recording, P=0.0822, Repeated measure One-way ANOVA Tukey’s test, n□=□7 neurons from 3 mice). **(M)** The box plot illustrates the average peak current (12-20 min) and summarize the findings on MOR-STD mechanism in the DLS (One-way ANOVA Dunnett’s multiple comparisons test). Time course data represent mean ± SEM. Box plots represent median and interquartile ranges and show average peak responses (12-20 minutes). ns=not significant, *P□<□0.05, **P□<□0.01.

### Presynaptic HCN1 channels are indispensable for MOR-mediated LTD

Given that we identified cAMP signaling as important for MOR-LTD expression (Fig. 6), we also explored the possibility of HCN1 channels as key factors for MOR-LTD. To do so, we applied the general HCN blocker ZD7288 (25 μM) to brain slices for at least 30 minutes before DAMGO application and during the entire recording. HCN inhibition blocked the expression of MOR-LTD (104 ± 17%; Fig. 10A-D), suggesting a novel role for HCN channels in DLS glutamatergic synaptic plasticity. Next, we characterized the effects of HCN channels in glutamatergic DLS synaptic transmission, by bath applying 25 μM ZD7288 for at least 15 minutes before recording sEPSCs. We found that the inhibition of HCN channels caused an enhancement in the frequency and a decrease in the amplitude of sEPSCs (Fig. 10E-G), without affecting rise and decay times (Fig. 10H and I). These data suggest, that HCN channels in the DLS both modulate synaptic transmission and mediate MOR-LTD.

**Figure 10.**
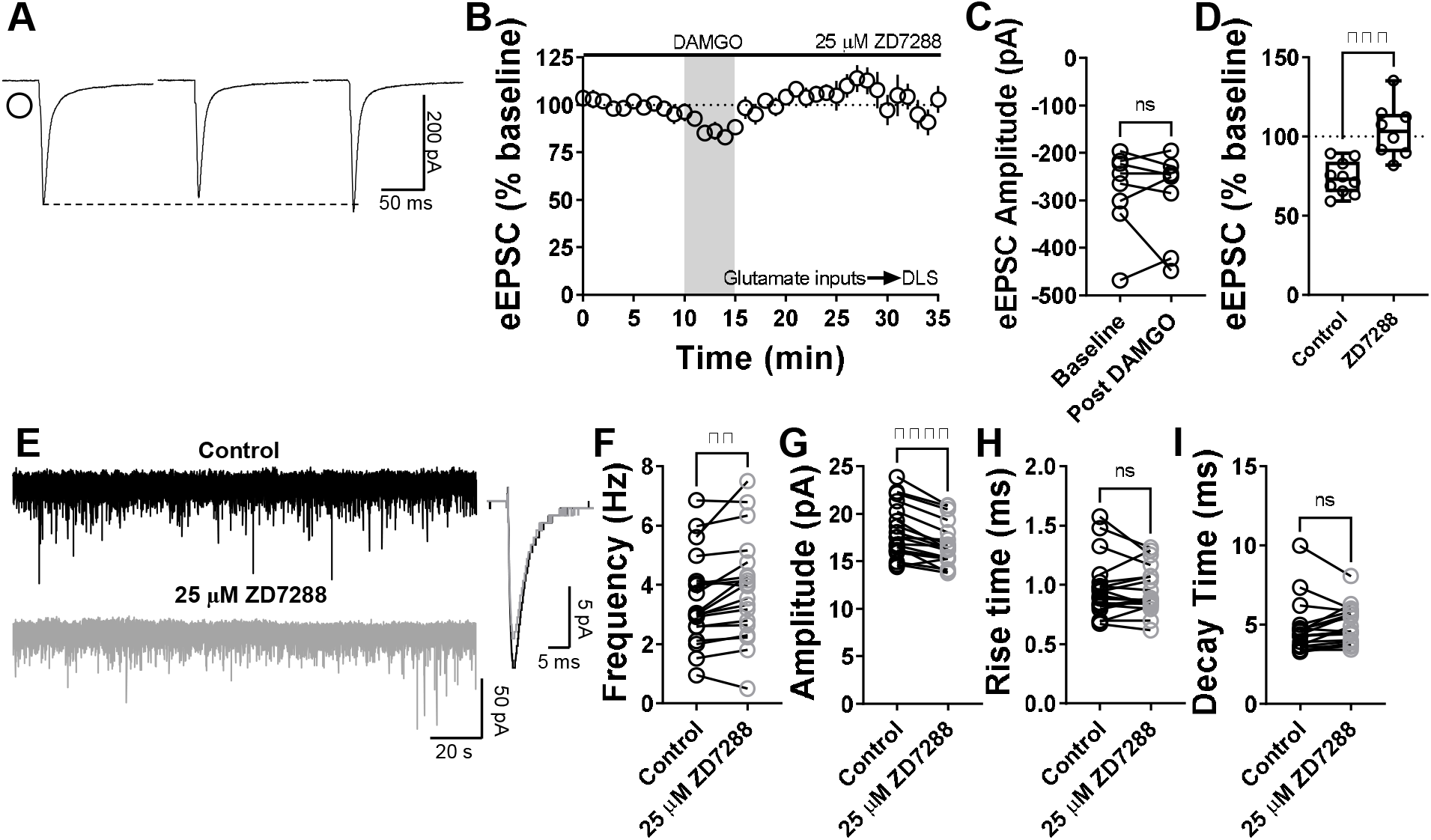
Inhibition of striatal HCN channels disrupts MOR-mediated LTD. **(A)** Representative eEPSC traces showing the effects of 25 μM ZD7288 before, during and after DAMGO (0.3□μM, 5□min) application. **(B-D)** HCN channel inhibition disrupted glutamatergic MOR-LTD (104 ± 6%, Unpaired t-test, P=0.000498, t16=4.67) in the DLS, showing no changes in eEPSC amplitude after DAMGO bath application (0–10□min baseline v. final 10□min of recording; paired t-test, P□=□0.635, t7□=0.496, n□= 8 neurons from 4 mice). **(E)** Representative synaptic current traces from C57BL/6J mice showing the spontaneous excitatory postsynaptic currents (sEPSC) in absence (black trace) or presence of 25 μM ZD7288 (gray trace, 15 min) application. **(F-I)** HCN channel inhibition increased frequency (paired t-test, P□=□0.00600, t18□=3.113, n□= 19 neurons from 3 mice) and decreased amplitude (paired t-test, P<0.0001, t18□=5.88, n□= 19 neurons from 3 mice), without changes in rise (paired t-test, P□=□0.608, t18□=0.522, n□= 19 neurons from 3 mice) and decay times (paired t-test, P□=□0.205, t18□=1.314, n□= 19 neurons from 3 mice) of the sEPSC events in DLS. Time course data represent mean ± SEM. Box plots show average of the final 10 minutes of recording and represent median and interquartile ranges. ns= not significant, **P□<□0.001, ***P□<□0.001, ****P□<□0.0001.

To further characterize this new role of HCN channels and specifically establish that presynaptic HCN1 channels are involved in the mechanism of expression of MOR-LTD, we used conditional HCN1 knockout mice (HCN1-flox) injected with AAV-cre vector into AIC to reduce the expression of HCN1 channel specifically in the AIC, and used AAV-GFP vector-infused mice as controls (Fig. 11A, B and Fig. 12). We used qPCR to confirm that AAV-cre infusion reduced HCN1 expression in the AIC (cre: 0.65 ± 0.2 vs GFP: 1.07 ± 0.4, Fig. 11C). Next, using electrical stimulation and recording in the DLS, we found that the ablation of HCN1 expression in the AIC reduced the magnitude of MOR-LTD in AAV-cre-injected mice, relative to AAV-GFP-injected mice (cre: 91 ± 12% vs GFP: 77 ± 10%; Fig. 11D-H). Next, we tested if the reduction of presynaptic HCN1 channels produced changes in basal striatal glutamatergic transmission. We found that the decrease of AIC HCN1 channels did not affect any synaptic parameters (Fig. 11I-M). These data suggest that HCN1 channels specifically on AIC inputs are necessary for MOR-LTD expression in the DLS.

**Figure 11.**
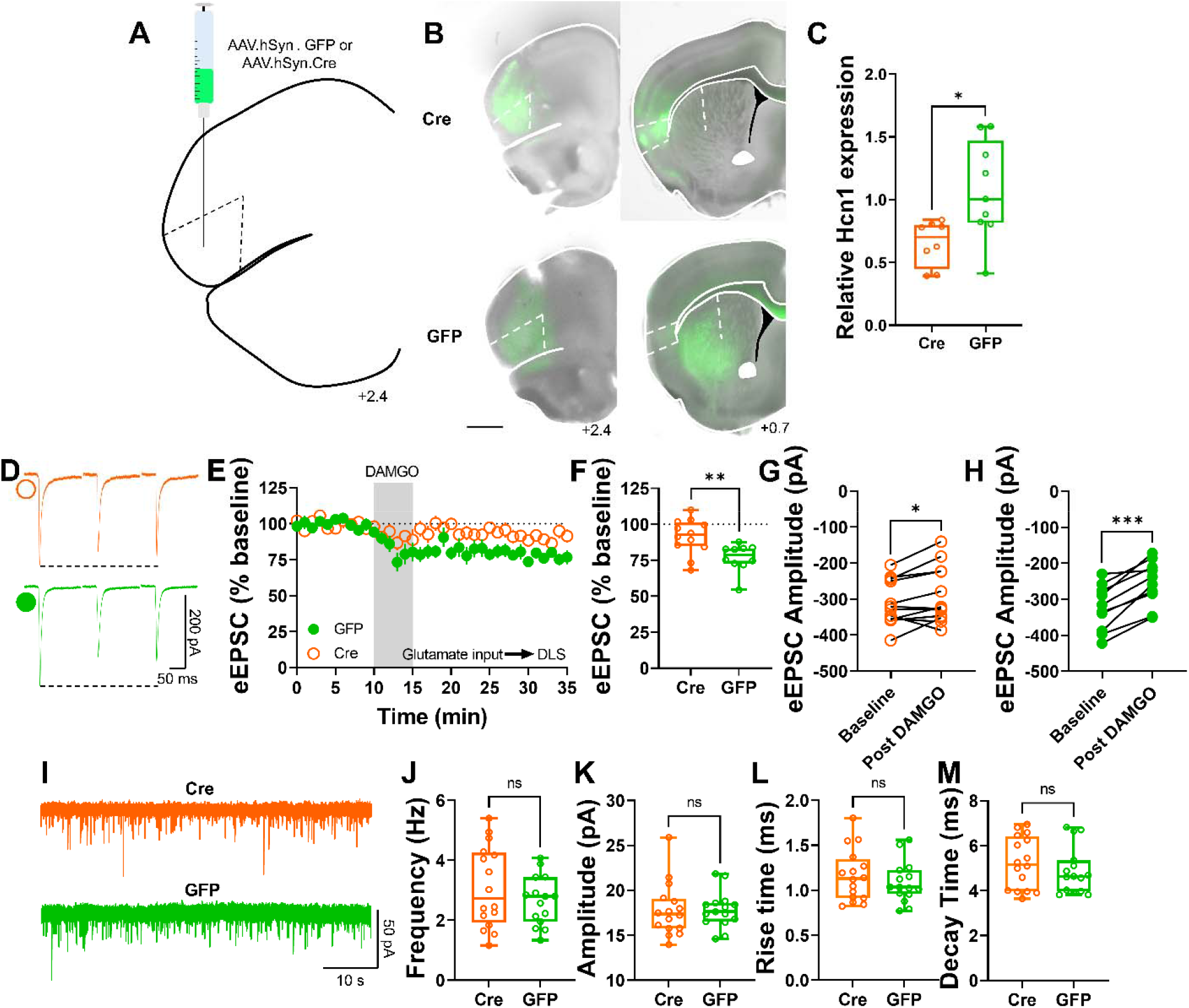
Genetic ablation of HCN1 channels in the AIC reduce MOR-mediated LTD expression in the DLS. **(A)** Schematic figure of coronal brain slice showing injection paradigm enabling knock out of HCN1 channel expression in AIC in HCN1-flox mice. An AAV vector encoding for either cre-recombinase (AAV.hSyn.cre) or eGFP (AAV.hSyn.eGFP) was injected at least 4 weeks prior to recordings. **(B)** Coronal brain slice showing the AAV-cre or AAV-GFP infection of AIC and dorsal striatal terminal expression (bar scale□=□1000□m). **(C)** AAV.cre reduced the expression of HCN1 in the AIC after 4 weeks post injection (cre: 0.65 ± 0.06 vs GFP: 1.07 ± 0.13, Unpaired t-test, P=0.0139, t15=2.785, n=8 cre-injected mice and n=8 GFP-injected mice). **(D)** Representative eEPSC traces showing the effects of DAMGO (0.3□μM, 5□min) application in brain slices of AAV-injected HCN1-flox mice. **(E-F)** The conditional decrease of HCN1 channel from AIC, blunted MOR-LTD in DLS (cre: 91 ± 4% vs GFP: 77 ± 3%, Unpaired t-test, P=0.00634, t20=3.05, n=12 neurons from 6 cre-injected mice and n=10 neurons from 4 GFP-injected mice). **(G)** HCN1 reduction did not fully disrupt MOR-LTD, because there was still a small, but significant reduction of eEPSC amplitude after DAMGO application (0–10□min baseline v. final 10□min of recording; paired t-test, P□=□0.0415, t11□=2.31, n□= 12 neurons from 6 mice). **(H)** Control GFP-injected mice showed normal decrease of eEPSC amplitude after DAMGO application 0–10□min baseline v. final 10□min of recording; paired t-test, P□=□0.000101, t9□=6.55, n□= 10 neurons from 4 mice). **(I)** Representative synaptic current traces from HCN1-flox mice showing the spontaneous excitatory postsynaptic currents (sEPSCs) in AAV-cre (orange trace) and AAV-GFP (green trace) injected mice. **(J-M)** HCN1 channel ablation from AIC synapses did not affect frequency (cre: 3.1 ± 0.3 Hz vs GFP: 2.7 ± 0.2 Hz, Unpaired t-test, P=0.277, t29=1.11, n=16 neurons from 4 cre-injected mice and n=15 neurons from 3 GFP-injected mice), amplitude (cre: 17.8 ± 0.7 pA vs GFP: 17.8 ± 0.5 pA, Unpaired t-test, P=0.980, t29=0.025, n=16 neurons from 4 cre-injected mice and n=15 neurons from 3 GFP-injected mice), rise time (cre: 1.16 ± 0.07 ms vs GFP: 1.09 ± 0.06 ms, Unpaired t-test, P=0.415, t29=0.822, n=16 neurons from 4 cre-injected mice and n=15 neurons from 3 GFP-injected mice) and decay time (cre: 5.24 ± 0.29 ms vs GFP: 4.9 ± 0.27 ms, Unpaired t-test, P=0.395, t29=0.396, n=16 neurons from 4 cre-injected mice and n=15 neurons from 3 GFP-injected mice) of the sEPSC events in DLS. Time course data represent mean ± SEM. Box plots show average of the final 10 minutes of recording and represent median and interquartile ranges. ns= not significant, *P□<□0.05, **P□<□0.01, ***P□<□0.001.

**Figure 12.**
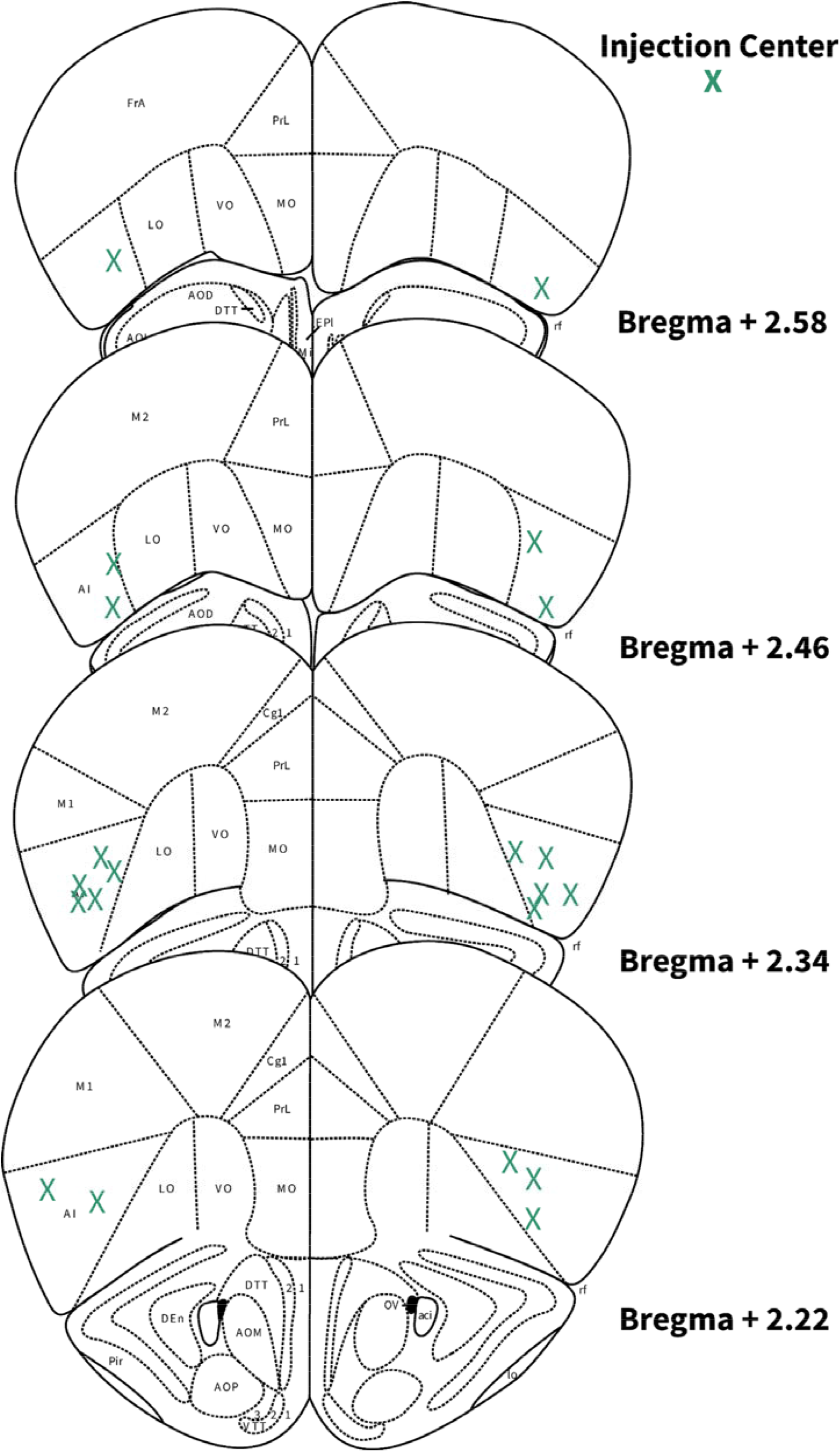
Locations of AIC injections for DLS electrophysiology. Schematic images of coronal brain slices modified from Allen Mouse Brain Atlas showing the bilateral injection centers of AAV-cre and EGFP in the AIC of HCN1-flox mice as verified by histology.

## Discussion

The current study demonstrates that the mechanism of MOR-mediated LTD is similar to the mechanism of induction of glutamatergic CB-LTD in that it involves cAMP-PKA-protein translation signaling and both occur at AIC-DLS synapses (Lonart *et al*., 2003; Yin *et al*., 2006; Mato *et al*., 2008; Yasuda *et al*., 2008; Atwood *et al*., 2014b; Younts *et al*., 2016). However, we found notable differences such as a lack of dependence on mTOR signaling (Fig. 8) and a novel mechanism involving HCN1 signaling for MOR-LTD (Fig. 10 and 11). Figure 13 summarizes the findings of the paper and illustrates the differential mechanisms involved in the expression of MOR-mediated LTD. Furthermore, the MOR-LTD mechanism is different than that of MOR-STD (Fig. 13), suggesting that MOR-mediated plasticity does not utilize a common pathway at all glutamatergic synapses.

**Figure 13.**
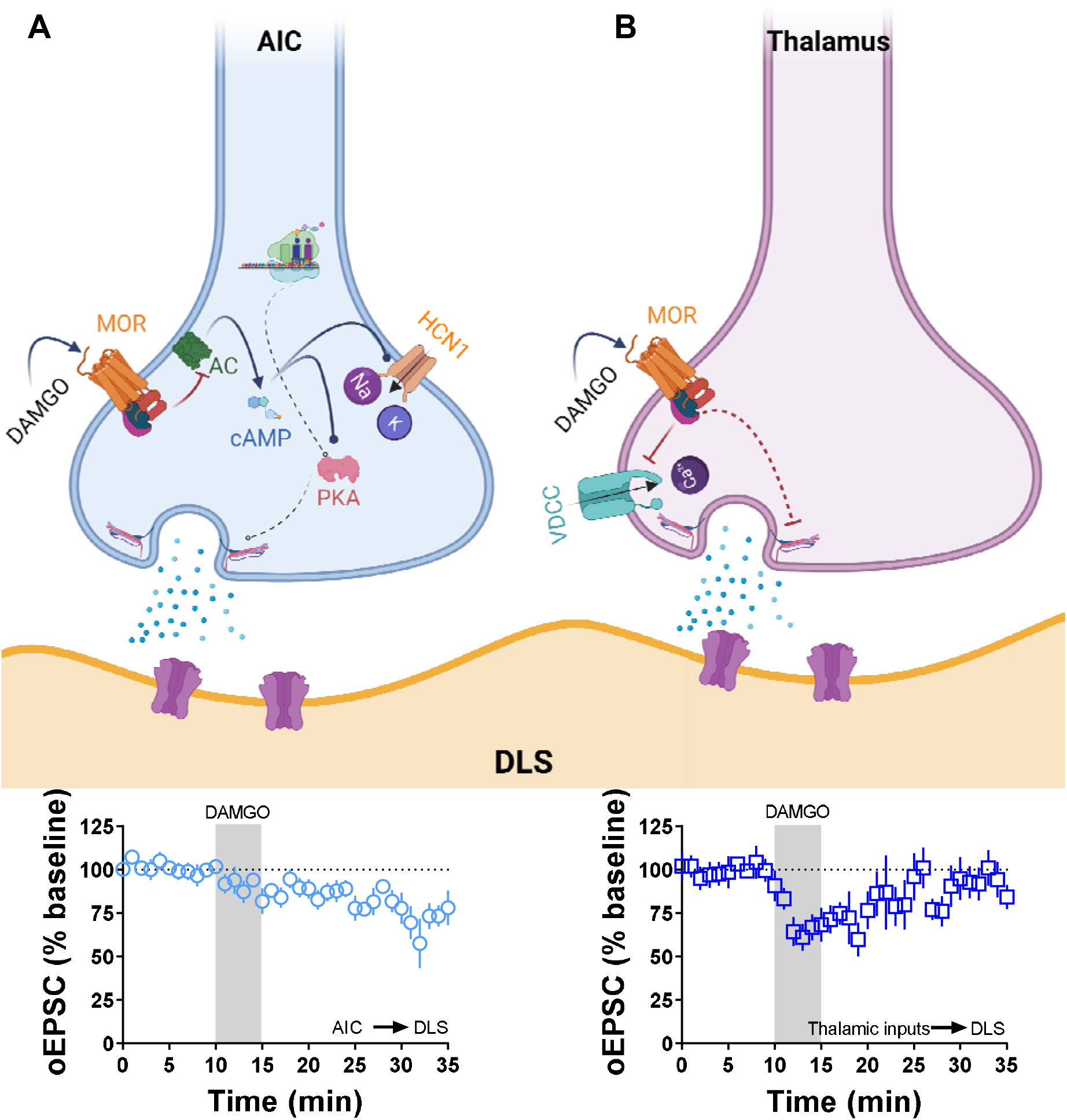
Summary of the proposed presynaptic signaling pathways that are involved in MOR-mediated LTD and STD. **(A)** The activation of MOR at AIC inputs in the DLS induces a decrease in the production of cAMP (via adenylyl cyclase (AC) inhibition), which impacts the activation of HCN1 channels and the function of presynaptic PKA. Furthermore, the presence of local protein synthesis is critical for the maintaining of the glutamatergic LTD shown below. **(B)** The activation of MOR at thalamic inputs in the DLS does not utilize the mechanisms illustrated in **(A)**. Therefore, the mechanism is conceptualized to involve MOR-driven Gβγ block of voltage dependent calcium channels (VDCC) producing a transitory reduction in the release of glutamate in the DLS as shown below. Schematic figure was created with BioRender.com.

Our previous report demonstrates that MORs expressed in AIC are the sole source of MOR-mediated LTD at glutamate inputs to the DLS(Munoz *et al*., 2018). However in the DMS, MORs on medial prefrontal cortex, anterior cingulate cortex, and basolateral amygdala inputs are capable of inducing LTD(Muñoz *et al*., 2020). MOR-LTD also occurs locally at cholinergic interneuron synapses that co-release glutamate on to MSNs(Munoz *et al*., 2018; Muñoz *et al*., 2020). However, alcohol only disrupts MOR-LTD at AIC inputs to DLS and at medial prefrontal and anterior cingulate cortex inputs to DMS leaving MOR plasticity at thalamostriatal, amygdalostriatal and cholinergic interneurons unaffected(Munoz *et al*., 2018; Muñoz *et al*., 2020). We also found that the opioid oxycodone disrupts MOR-LTD in DLS(Atwood *et al*., 2014a), and given that this is mediated by AIC-DLS MORs(Munoz *et al*., 2018), it follows that alcohol and opioids have similar actions on MOR plasticity at this synapse. Given that alcohol (and presumably oxycodone and other opioids) do not affect MOR plasticity at these other non-corticostriatal synapses, differential mechanisms that underlie MOR plasticity at each type of synapse may confer susceptibility to the deleterious effects of these drugs and could account for their mechanisms of action in the development of habitual or compulsive use of the drugs.

Previous work has shown that the inhibition of PKA disrupts CB-LTD in the nucleus accumbens(Mato *et al*., 2008) and that presynaptic PKA is required to induce cannabinoid LTD in cerebellum(Lonart *et al*., 2003) and cannabinoid synaptic plasticity in hippocampus(Yasuda *et al*., 2008). Moreover, previous reports had shown that the activation of AC by forskolin enhances corticostriatal EPSC amplitudes(Cho *et al*., 2008) and blocked delta opioid receptor-mediated LTD of inhibitory postsynaptic currents(Patton *et al*., 2016). cAMP/PKA signaling has also been implicated in presynaptic type 2/3 metabotropic glutamate receptor- and type 1b serotonin receptor-mediated LTD at various synapses in the brain(Atwood *et al*., 2014a). As previously reported(Duffy & Nguyen, 2003), we confirmed that the activation of postsynaptic PKA is necessary for the induction of LTP in CA1 pyramidal neurons from the hippocampus, but not for MOR-LTD in DLS (Fig. 5). Until now, the role of cAMP/PKA signaling pathway in MOR-mediated plasticity had not been addressed. Our results demonstrate that presynaptic PKA inhibition blocked AIC-DLS MOR-LTD (Fig. 4 and 13), but not thalamostriatal MOR-STD (Fig. 9D-F and 13). Interestingly, the magnitudes and time courses of eEPSCs and oEPSCs did not differ between measures (Fig. 4A, D, H, K, O and R), suggesting that PKA is not required for an acute MOR-mediated inhibition of glutamate release, but it is required for the maintenance of MOR-LTD. On the other hand, we found that AC activation disrupted both acute MOR-mediated inhibition (in fact turning it into a transient potentiation) and MOR-LTD (Fig. 6 and 13), but had no effect on MOR-STD (Fig. 9G-I and M and 13). Altogether these data suggest that cAMP/PKA pathway may be a general mechanism whereby GPCRs that signal through Gαi may induce LTD, but a receptor’s ability to couple to Gαi does not necessarily mean that it will induce cAMP/PKA-dependent LTD upon activation.

Protein translation is necessary for striatal CB-LTD(Yin *et al*., 2006) and is involved in GPCR-mediated LTD at some synapses, but not others(Atwood *et al*., 2014a; Younts *et al*., 2016) suggesting that protein translation can be an important component of glutamatergic LTD, depending on the synapse. Indeed, local protein translation can take place in the presynaptic region, where the activation of CB1 receptors enhances protein translation, being critical for the induction and not the maintenance of inhibitory LTD(Younts *et al*., 2016). Here we demonstrated that protein translation is necessary to produce MOR-LTD (Fig. 7 and 13), but not MOR-STD (Figure 9J-M and 13). Our data is consistent with the previous reports on protein translation-dependent CB-LTD, and suggest that it is presynaptic protein translation that is involved in MOR-LTD expression. However, cycloheximide has been reported to also activate p38/MAPK signaling (Lockhead *et al*., 2020) and several studies have shown a critical role of p38 kinases in the induction of LTD and LTP (Asih *et al*., 2020). Therefore, more data is needed to determine this is definitively protein translation-mediated in the DLS.

Although we have previously reported that MOR- and CB-LTD are mutually occlusive in the DLS(Atwood *et al*., 2014a), and here we found that there were similarities between MOR-LTD and CB-LTD mechanisms, we found some differences which make both types of LTD distinct. In the DLS, MOR-LTD of glutamate input is restricted to the AIC(Munoz *et al*., 2018), which differs from the more broadly expressed (and overlapping at AIC-DLS synapses, Fig. 3) CB-LTD(Wu *et al*., 2015; Gremel *et al*., 2016), suggesting that these two forms of LTD may occur in distinct synaptic environments that could utilize differing mechanisms. It is likely that even in synapses where both receptors are expressed (e.g. in AIC terminals) that each receptor shares partially overlapping, but not identical signaling pathways.

A clear example of distinct signaling mechanisms identified here is in the role of the mTOR pathway in CB-LTD and MOR-LTD. The CB1 receptor can signal via mTOR in the hippocampus, where the increase of protein translation induced by CB1 receptor activation is mTOR-dependent producing inhibitory CB-LTD(Younts *et al*., 2016). We predicted that we would see the same effect in the DLS for both CB-LTD and for MOR-LTD. Partially correct in our prediction, we found that CB-LTD is blocked after the inhibition of mTOR in the DLS (Fig. 8A-D), but the mechanism of MOR-LTD did not utilize mTOR (Fig. 8E-H). These novel data regarding CB1-mTOR signaling in DLS provide rationale for future experiments addressing the role of CB1-mTOR signaling in dorsal striatal-related behaviors.

Presynaptic cations such as Na^+^ and K^+^ are important for the propagation of action potentials and subsequent glutamate release(Chen & Lui, 2021). HCNs channels are permeable to Na^+^ and K^+^ and modulated by cAMP(Sartiani *et al*., 2017), and they can decrease neurotransmission by restricting presynaptic Ca^2+^ influx(Huang *et al*., 2011). We did identify a critical role for the HCN1 channel in MOR-LTD expression (Fig. 10, 11 and 13). It has been reported that HCN1 channels can be found at presynaptic sites in the globus pallidus, where the inhibition by ZD7288 produced an increase of miniature inhibitory postsynaptic current frequency, with no changes in amplitude(Boyes *et al*., 2007). The block or deletion of HCN1 causes an increase in the frequency of miniature EPSCs in entorhinal cortical pyramidal neurons(Huang *et al*., 2011). Here we showed that broad blockade of HCN channels alters glutamatergic synaptic transmission release (Fig. 10E-I). However, when we deleted just HCN1 from AIC inputs to DLS (Fig. 11C), we did not observe any changes in basal glutamate transmission (Fig. 11I-M). The differences in effects of broad blockade and synapse-specific deletion may be attributed to the overall contribution of HCN1s at AIC inputs relative to the population of glutamate synapses. Genetic deletion did not completely ablate MOR-LTD expression (Fig. 11G). This could be due to an incomplete AAV vector transduction (most recordings had a complete loss of LTD) or perhaps that HCN1 contributes to LTD differently at AIC inputs to different subtypes of MSNs. Therefore, in the future, it will also be important to investigate the role of HCN1 in MOR-LTD at AIC inputs to D1- or D2-expressing MSNs (direct or indirect pathway respectively). It has recently been reported that HCN1 contributes to alcohol preference(Salling & Harrison, 2020). Given that alcohol disrupts MOR-LTD in DLS, this mechanism of MOR-HCN1 signaling may be a target of alcohol’s effects and contribute to alcohol-related, dorsal striatal-dependent behaviors such as habitual and compulsive alcohol drinking. As the opioid oxycodone also disrupts MOR-LTD it will be intriguing to determine the impact of both oxycodone and alcohol on AIC-DLS HCN1 function. HCN1 may therefore have therapeutic potential in alcohol and opioid use disorders.

In conclusion, these findings indicate that glutamatergic synapses in the DLS have distinct mechanisms of plasticity induction mediated by the same types of receptors. The specific signaling processes that result in these different forms of plasticity might be the answer to why different types of plasticity are differentially sensitive to drugs of abuse and alcohol (Fig. 13) (Atwood *et al*., 2014b; Munoz *et al*., 2018; Muñoz *et al*., 2020). However, further work is required to resolve what components of these mechanisms render some synapses susceptible to the deleterious effects of these drugs while others are resistant. It will also be important to decipher the behavioral relevance of AIC-DLS synapse HCN1 channels and CB1-mTOR signaling in the DLS.

## Additional Information

### Data availability statement

All data in this study are available upon reasonable request to the corresponding author. All reagents and other materials used for this work are commercially available.

### Author Contributions

B.M., and B.K.A., designed experiments, discussed the results, contributed to all stages of manuscript preparation and editing. BM performed and analyzed all experiments. B.M.F., assisted in collecting electrophysiological data. F.Y. performed qPCR experiment. All authors contributed to and approved the final version of the manuscript.

## Supporting information

Additional supporting information can be found online in the Supporting Information section at the end of the HTML view of the article. Supporting information files available:

Peer Review History

Statistical Summary Document

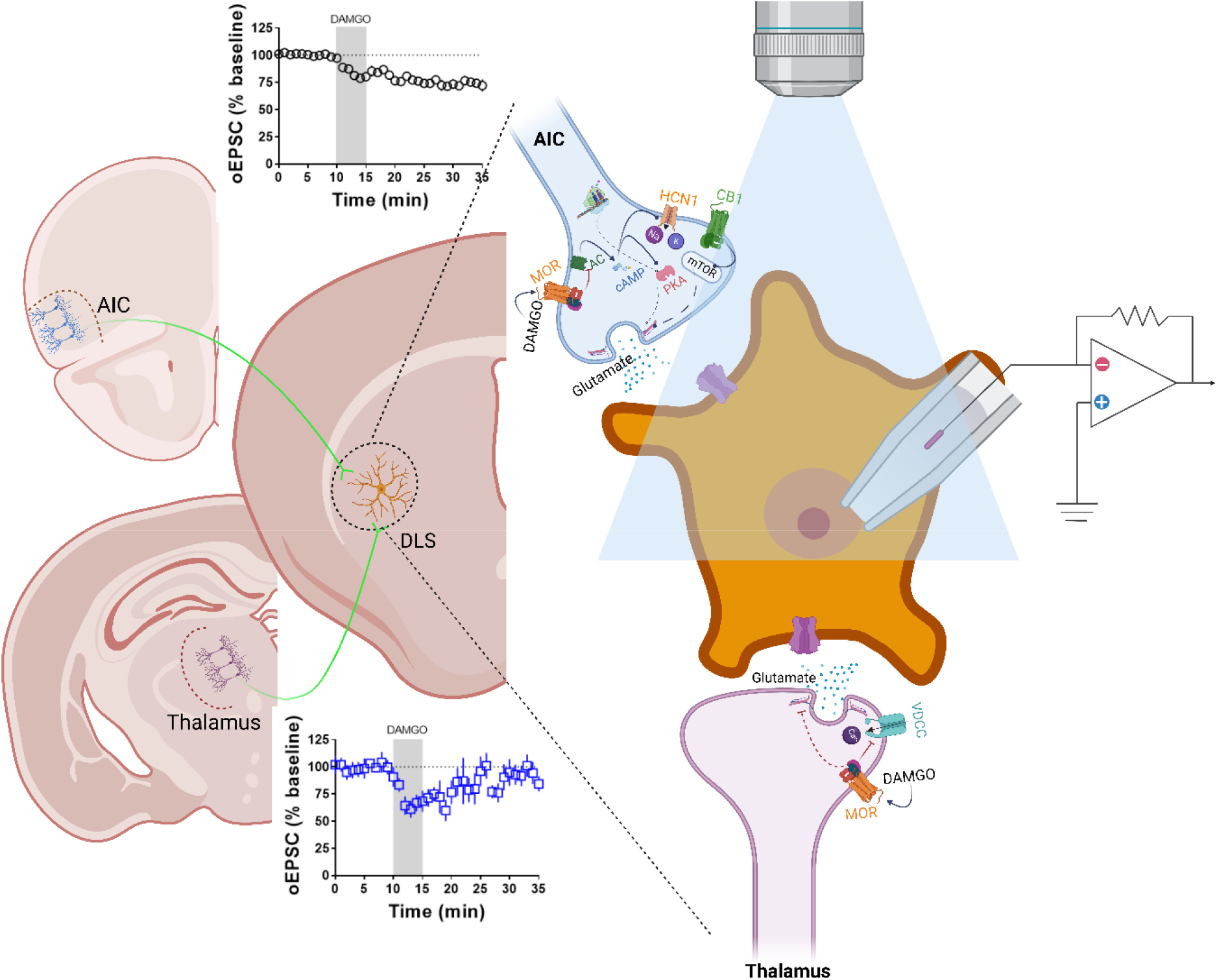

## Graphical Abstract Figure Legend

We tested the hypothesis that mu opioid receptor (MOR) plasticity utilizes different mechanisms at distinct glutamatergic synapses in the dorsolateral striatum (DLS). Glutamatergic synapses from the anterior insular cortex (AIC) expressed MOR-mediated long-term depression (MOR-LTD), and synapses from the thalamus expressed MOR-mediated short-term depression (MOR-STD). Using brain slice patch-clamp electrophysiology combined with optogenetics and pharmacology, we found that MOR from AIC inputs required different mechanism to induce LTD, than MOR from thalamic inputs. Interestingly, MOR-LTD shares similar mechanisms with cannabinoid-LTD (CB-LTD). However, MOR-LTD does not require mTOR signaling, while CB-LTD does. We characterized the role of presynaptic HCN1 channels in MOR-LTD induction as HCN1 channels expressed in AIC are necessary for MOR-LTD expression in the DLS. These results suggest a mechanism in which MOR activation needs HCN1 to induce MOR-LTD, suggesting a new target for pharmacological modulation of synaptic plasticity, providing new opportunities to develop novel drugs to treat alcohol and opioid use disorders.

